# Tumor-infiltrating CD27^-^IgD^-^ regulatory B cells suppress cytotoxic CD8^+^T cell responses in renal cell carcinoma

**DOI:** 10.1101/2025.07.08.663720

**Authors:** Zara Baig, Isabella Withnell, Joseph C.F. Ng, Christopher J.M. Piper, Hannah F. Bradford, Antonio Rullan, Edward H. Arbe-Barnes, Camila G.X. de Brito, Alessandro Di Tullio, Thomas J. Mitchell, Hans J. Stauss, Maxine G.B. Tran, Franca Fraternali, Claudia Mauri

## Abstract

B cells play a pivotal role in shaping the tumor microenvironment (TME) and tertiary lymphoid structures (TLS), but the functions of specific B cell subsets in cancer pathogenesis remain unclear. Using a novel tissue-centric single cell RNA-sequencing (scRNA-seq) bioinformatic workflow aimed at unraveling cancer-specific clusters, combined with flow cytometry and high-plex spatial imaging, we identify a renal cell carcinoma (RCC)-specific enrichment of CD27^-^IgD^-^CD21^+^CD11c^-^ double negative 1 (DN1) B cells associated with worse prognosis and regulatory function. Spatial profiling localizes DN1 Bregs within immature TLSs, in close proximity to IL-10⁺ and TGFβ⁺CD8⁺T cells. RCC-resident DN1 B cells are enriched for endosomal Toll-like receptor (TLR) signaling pathways and stimulation of tumor-infiltrating lymphocytes (TILs) with TLR agonists induces the differentiation of IL-10^+^ and TGFβ^+^DN Bregs suppressing CD8^+^T cell cytotoxicity, partially via IL-10 and TGFβ. These findings identify a pro-tumorigenic B cell population with potential diagnostic and therapeutic relevance in RCC.

## Introduction

B cells play a crucial role in shaping local immune responses across different tissue microenvironments. While more is known about the phenotype and function of circulating B cell subsets, comparatively less is known about tissue-resident B cells and how their phenotype and function are shaped by the tumor microenvironment (TME). Tumor-infiltrating B cells (TIBs) within tertiary lymphoid structures (TLSs) express high levels of MHC class I and II molecules, facilitating antigen presentation to T cells with cytotoxic activity^1–4^. Additionally, antibody-producing plasma cells (PCs) can mediate tumor cell destruction through antibody-dependent cellular cytotoxicity (ADCC) and complement activation^5–8^. B cells produce both pro- and anti-inflammatory cytokines, further regulating immune-tumor responses^9^. The latter function has been postulated to be exerted by regulatory B cells (Bregs), an immunosuppressive subset that may promote tumor progression by secreting pro-tumorigenic cytokines, including IL-10 and TGFβ, inhibiting cytotoxic T cells and inducing regulatory T cells (Tregs)^10–14^. The maturity and cellular constitution of TLSs have been linked to prognosis outcome, with mature TLSs generally associating with favorable prognosis across many solid cancers, including lung, ovarian, melanoma, and head and neck cancers^15–18^.

Pan-cancer atlases have begun to elucidate the diverse functional states of TIBs, distinguishing between putative pro- and anti-tumorigenic subsets that differentially influence patient outcomes across cancer types^19–21^. While these studies offer detailed analyses of B cell subset composition, they are often dominated by cancers with high B cell infiltration, potentially undermining functionally relevant distinctions in cancers with fewer B cells. One example is renal cell carcinoma (RCC), which exhibits lower B cell infiltration compared to other cancer types such as colon and lung cancer^20^. RCC represents a group of kidney cancers originating from the epithelial cells of the nephron in the renal cortex^22^. RCC accounts for approximately 2% of all adult cancers, with clear cell RCC (ccRCC) being the most common subtype, representing 75-80% of cases^23,24^. RCC mortality remains a global health concern, with over 140,000 deaths reported annually^24^. While early detection greatly improves survival, it is often incidental, and advanced-stage RCC is associated with significantly worse survival^23^. Here we provide a comprehensive characterization of B cells in RCC, identifying a population of DN1 Breg enriched in the tumor and tumor margin compared to matching blood and background kidney (BK). *In vitro*, we show that DN B cells suppress IFNγ expressing CD8^+^T cells differentiation. Together, our work offers implications for better understanding immune escape mechanisms and designing B cell subset-specific, targeted immunotherapies.

## Results

### Independent clustering of B cells across cancer types reveals an enrichment of DN B cells in RCC

To investigate how different tissue microenvironments and varying levels of B cell infiltration drive the differentiation of resident B cells, we compared four cancer types, breast carcinoma (BRCA), colon adenocarcinoma (COAD), lung cancer (LC), and RCC, representative of mucosal and non-mucosal tissues. To preserve cancer-driven clusters, we applied single-cell Variational Inference (scVI) to separately integrate and cluster the available single-cell B cell datasets for each cancer type (Table S1A, Figure S1A for schematic describing the pipeline). In large-scale benchmarks, scVI consistently ranks among the top integration methods, correcting both technical and biological variability, and generates a denoised expression matrix that enhances detection of low abundance genes^25^. After filtering out low-quality cells and excluding plasma cells to preserve better B cell subset granularity, a total of 25,427 single-cell B cell transcriptomes were obtained from 13 BRCA patients, 28,054 from 15 COAD patients, 26,652 from 15 LC patients, and 18,782 from 9 RCC patients (Table S1A).

Clustering of the scVI-derived embeddings identifies 14 transcriptionally distinct TIB clusters in BRCA, 13 in COAD, 14 in LC, and 15 in RCC (Figure 1A). We trained an additional scVI model to project B cells from each cancer into a shared latent space to identify cancer-specific and shared clusters (Figure S1A). To quantify transcriptomic similarity, we calculated cosine distances between the centroids of each cluster. Low cosine distances between nearby clusters in the shared latent space reflect similarities in underlying gene expression (Figure 1B). Hierarchical clustering based on the cosine distances organized B cells into different groups (Figure 1B). We then evaluated known B cell marker genes to validate these groups and define their biological identity.

**Figure 1.**
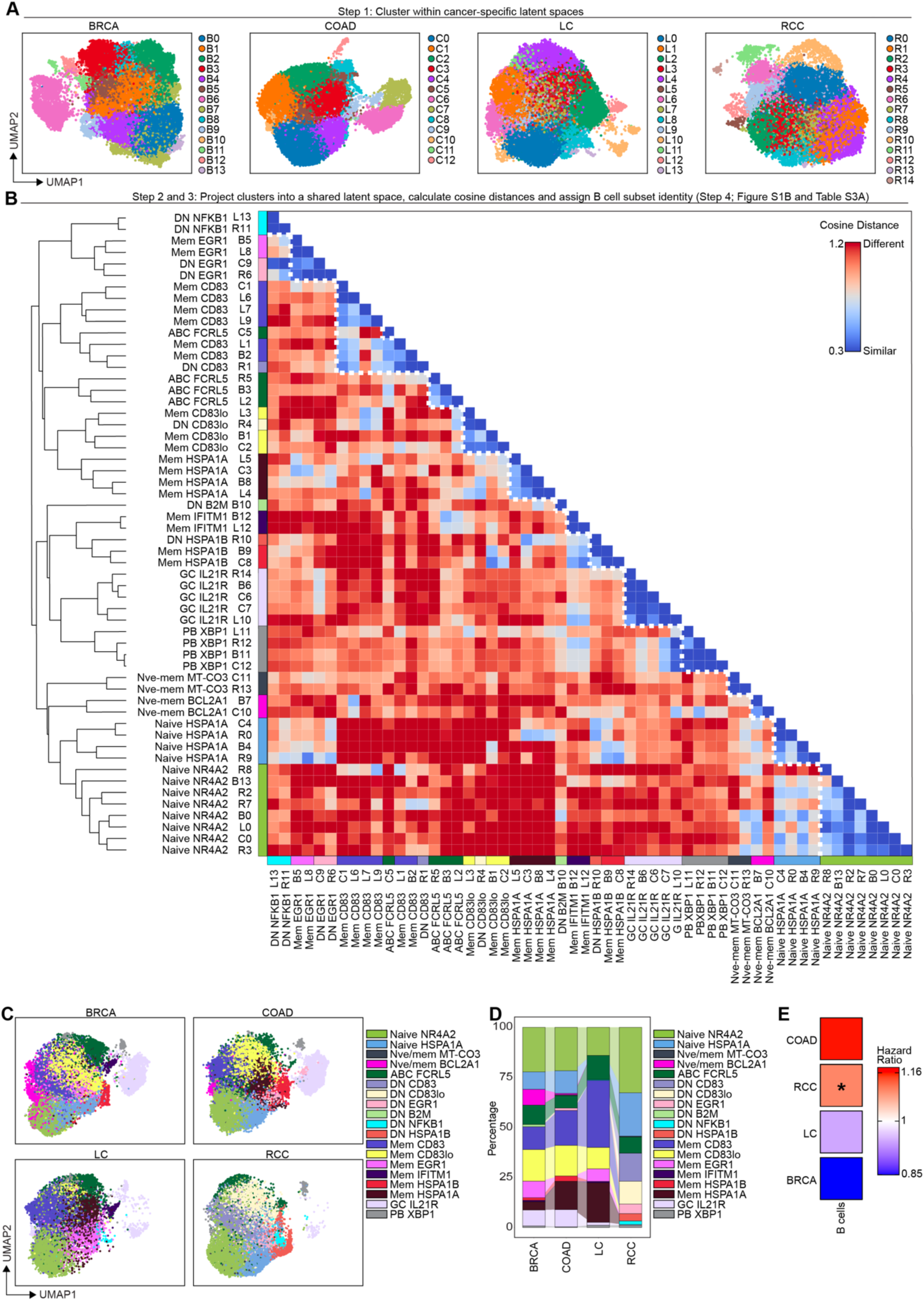
Tissue-driven clustering of B cells, followed by projection into a shared latent space, reveals an expansion of DN B cells primarily in RCC. (A) Uniform manifold approximation and projection (UMAP) visualization of B cell clusters within the cancer-specific latent space for breast carcinoma (BRCA), colon adenocarcinoma (COAD), lung cancer (LC), and renal cell carcinoma (RCC). (B) Dendrogram and heatmap showing hierarchical clustering of transcriptionally similar cancer-specific clusters grouped according to cosine distance, annotated according to key gene markers (Figure S1B). DN = double negative, mem = memory, ABC = atypical B cell, GC = germinal center, PB = plasmablast, Nve = naive. (C) UMAPs showing annotated cancer-specific B cell clusters projected into the shared latent space. (D) Stacked bar chart showing the proportion of annotated B cell clusters in BRCA, COAD, LC, and RCC. (E) Prognostic association of B cell abundance with survival in BRCA, COAD, LC, and RCC estimated by CIBERSORT using The Cancer Genome Atlas (TCGA) data; \**p*<0.05. *p* values were derived by Cox proportional hazard model for each cancer type.

We assigned identities to each cluster according to known B cell maturation, function, and differential gene expression (DEG) (Figure S1B and Table S2). These include two *IGHD^hi^TCL1A^hi^*naive (naive *NR4A2,* naive *HSPA1A*), two *IGHD^lo^FCER2^hi^TNFRSF13B^hi^*naive-memory intermediate (naive-mem *BCL2A1*; naive-mem *MT-CO3*), one *ZEB2^hi^FCRL5^hi^* atypical memory B cell (ABC) (ABC *ITGAX*), six *CD27^lo^IGHD^lo^* DN (DN *NFKB1*; DN *EGR1*; DN *CD83*; DN *CD83lo*; DN *B2M*; DN *HSPA1A*), six *CD27^hi^CD38^lo^* memory (mem *EGR1*; mem *CD83*; mem *CD83lo*; mem *HSPA1A*; mem *IFITM1*; mem *HSPA1B*), a *SUGCT^hi^* GC *IL21R*, and a *PCNA^hi^* plasmablast (PB) *XBP1* population (Figure 1C and Figure S1B).

Identifying CD27^-^IGHD^-^ DN B cells by scRNA-seq is challenging, as it relies on relative gene expression of *CD27* and *IGHD* across clusters, which is less discriminative than measuring surface protein expression, for example, by flow cytometry. To aid the annotation of cancer-specific DN B cell clusters, we trained an additional scVI model and compare with the denoised expression of *CD27* and *IGHD* of naive, DN, and memory B cells purified from healthy peripheral blood^26^ (Table S3A). This analysis reveals that DN B cells constitute 35.7% of TIBs in RCC, compared to 1.1% in BRCA, 1.16% in COAD, and 0.36% in LC. These findings confirm that RCC harbors more DN B cells compared to the other cancers analyzed (Table S3B).

B cell subset proportions vary between cancers, with some clusters shared (e.g., ABC *FCRL5*) with others restricted to specific cancers (e.g., DN *HSPA1B* in RCC) (Figure 1D). COAD and LC exhibit an enrichment of memory *HSPA1A* B cells compared to BRCA and RCC, whereas GC *IL21R* B cells are enriched in BRCA and COAD but are scarce in LC and virtually absent in RCC. RCC displays a unique expansion of DN B cell clusters enriched in activation markers (*CD83, CD69, and NR4A2*) (Figure 1D and S1B), and an enrichment of naive *HSPA1A* B cells which shares multiple gene expression features with DN B cell populations (including heat shock genes *HSPA1A* and *HSPA1B,* and *EGR1)* (Figure S1B). Trajectory analysis, conducted in the individual cancer latent spaces to improve accuracy, infers that RCC-resident naive *HSPA1A* B cells give rise to DN B cells (e.g., DN *EGR1* B cells) (Figure S1C and S1D).

Compared to our tissue-driven bioinformatic pipeline, joint integration and clustering of B cells from BRCA, COAD, LC, and RCC reveals 14 shared clusters, many of which are broadly defined as memory or naive rather than DN B cells, highlighting a loss of granularity (Figure S2A-S2C). One example is the loss of the diverse RCC-resident DN populations, forming one DN *EGR1* cluster (Figure S2D). Prognostic associations further highlight the context dependence inter-tumoral B cell heterogeneity: high B cell abundance is significantly associated with poorer survival in RCC whilst linked to trends toward improved survival in LC and BRCA (Figure 1E).

### DN1 and DN3 B cell subsets are enriched in RCC

To determine whether the enrichment of DN B cells is kidney- or RCC-specific, next we took advantage of our in-house scRNA-seq dataset comprising matching tumor (*n*=147,917 single cells), tumor margin (*n*=20,534), adjacent BK tissue (*n*=11,201), and blood (*n*=52,914) from 10 treatment-naive RCC patients (Table S1B). As previously reported^27^, we identify 12 main cell lineages across tissues, based on the expression of canonical cell type markers (Figure S3A and S3B).

Unsupervised clustering of B cells and PCs across tissue compartments identifies 15 distinct populations (Figure 2A-2C). Proportionally, RCC blood contains higher frequencies of naïve and proliferative naive (*CD27^lo^IGHD^hi^TCL1A^hi^)* followed by resting DN1 (*CD27^lo^IGHD^lo^CD83^lo^)* and memory B cells (*CD27^hi^IGHD^lo^TNFRSF13B^hi^)*, including both classical and proliferative subsets, compared to other tissues (Figure 2B-2D). ABCs (*ZEB2^hi^TBX21^hi^FCRL5^hi^*) are also mainly present in blood. In both tumor and tumor margin tissues, DN1 B cells (*CD27^lo^IGHD^lo^CR2^hi^ITGAX^lo^)* were the predominant population amongst all B cells, followed by plasma cells (*PRDM1^hi^XBP1^hi^*), DN3 B cells (*CD27^lo^IGHD^lo^CR2^lo^ITGAX^lo^*), and a naive B cell population expressing stress-related genes (*CD27^lo^IGHD^mid^HSPA1A^hi^HSPA1B^hi^*), compared to blood and BK tissue. The BK is largely characterized by an activated kidney-resident B cell population, lacking the expression of several differentiation-stage B cell markers (*CD27^lo^IGHD^lo^CD38^lo^PCNA^lo^CD69^hi^BCL2A1^hi^*) (Figures 2B-2D).

**Figure 2.**
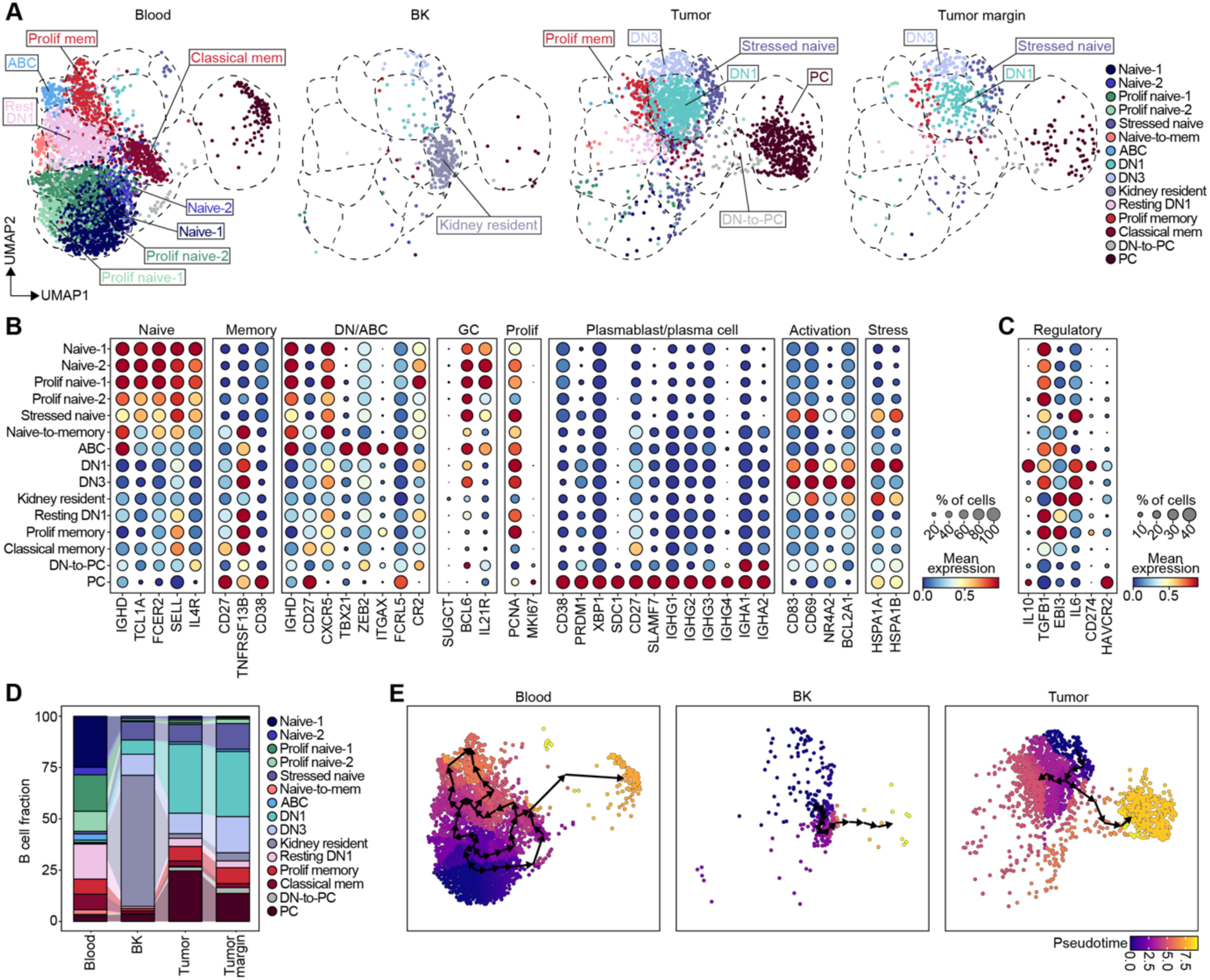
DN1 B cells are enriched in RCC and are predicted to arise from RCC-resident naive B cells. (A) UMAPs showing annotated B cell clusters in matching tumor, tumor margin, BK, and blood from 10 diagnostically confirmed RCC treatment-naive patients. Prolif = proliferation; naive-to-mem = naive-to-memory, rest = resting, mem = memory, DN1 = double negative 1, DN3 = double negative 3, DN-to-PC = double negative-to-plasma cell, PC = plasma cell. (B) Dotplot showing for each B cell cluster across paired tissues the proportion of cells expressing selected key B cell markers with scVI-denoised expression > 0.1 (point size) and their mean expression levels (color intensity), min-max scaled across annotated B cell clusters. (C) Dotplot showing for each B cell cluster across paired tissues the proportion of cells expressing selected key regulatory markers with scVI-denoised expression > 0.1 (point size) and their mean expression levels (color intensity), min-max scaled across annotated B cell clusters. (D) Stacked bar chart showing the proportion of annotated B cell clusters in blood, BK, tumor, and tumor margin. (E) UMAPs showing the inferred pseudotime trajectories of B cells in the blood, BK, and tumor. Arrows indicate the direction of trajectories.

Pseudotime trajectory analysis infers an extrafollicular pathway for tumor-associated B cells in contrast to their blood-resident counterparts. While naive B cells in the blood follow a canonical maturation trajectory (progressing from naive to memory B cells and to plasma cells), tumor-infiltrating naive B cells progress into DN1 B cells, followed by a DN-to-PC intermediate (*CD27^-^IGHA1^+^*), and into PCs (Figure 2E).

BCR-sequencing (data obtained from pan-cancer study^21^) shows that DN B cells generally exhibit lower levels of somatic hypermutation (SHM) in their *V_H_* genes compared to memory B cells (Figure S3C)^28^. With the exception of *CD83^lo^*DN, RCC DN B cells display lower SHM levels when compared to DN and memory B cells in COAD and LC (Figure S3C); RCC-resident plasma cells (annotated in a previous study^21^) also exhibit the lowest SHM levels among 15 cancer types analyzed (Figure S3D). Isotype analysis shows that approximately one-third of DN1 and DN3 B cells express IgM, while the remainder have undergone class switch recombination (CSR), predominantly to IgA and to a lesser extent IgG (Figure S3E).

### DN1 B cells enriched in freshly explanted RCC tissue are associated to poor prognosis

To corroborate our single-cell transcriptomic results at protein level, we evaluated B cell subset composition by flow cytometry in an external cohort of 34 paired tumor and BK tissues from treatment-naive RCC patients undergoing radical nephrectomy (Table S4A). We were unable to collect tumor margin specimens as this region is required by the pathologist to stage disease severity. Normalization per gram of tissue reveals significantly higher CD45^+^lymphocyte infiltration, including CD19^+^B cells and CD3^+^T cells, in tumors compared to BK (gating strategy in S4A, Figure S4B-F). Tumors show a significant increase in DN B cells (CD27^-^IgD^-^) and plasma cells (CD24^lo/-^ CD38^hi^BLIMP-1^+^), mirrored by a significant reduction in CD27^+^IgD^-^B cells, unswitched memory (USM; CD27^+^IgD^+^), and class-switched memory (CSM; CD27^+^IgD^-^ CD24^+^CD38^-^IgM^-^IgD^-^) subsets (Figure 3A-3C). No significant differences between tumor and BK are observed for transitional (CD27^-^IgD^+^CD24^hi^CD38^hi^), mature naive (CD27^-^IgD^+^CD24^int^CD38^int^), or IgM^+^ memory (CD27^+^IgD^-^CD24^+^CD38^-^IgM^+^IgD^-^) B cells (Figure 3C and 3D). Within the DN B cell population, we confirm a significant increase in DN1 (CD21^+^CD11c^-^) B cells, while no significant differences are observed for DN2 (CD21^-^CD11c^+^), DN3 (CD21^-^CD11c^-^), or DN4 (CD21^+^CD11c^+^) B cell subsets in the tumor compared to BK (Figure 3E). Although CXCR5 has been previously used to identify DN1 B cells in the blood^29^, CD21 is used in lieu of CXCR5 since the latter could be upregulated on B cells in response to CXCL13 produced by RCC cells^30^.

**Figure 3.**
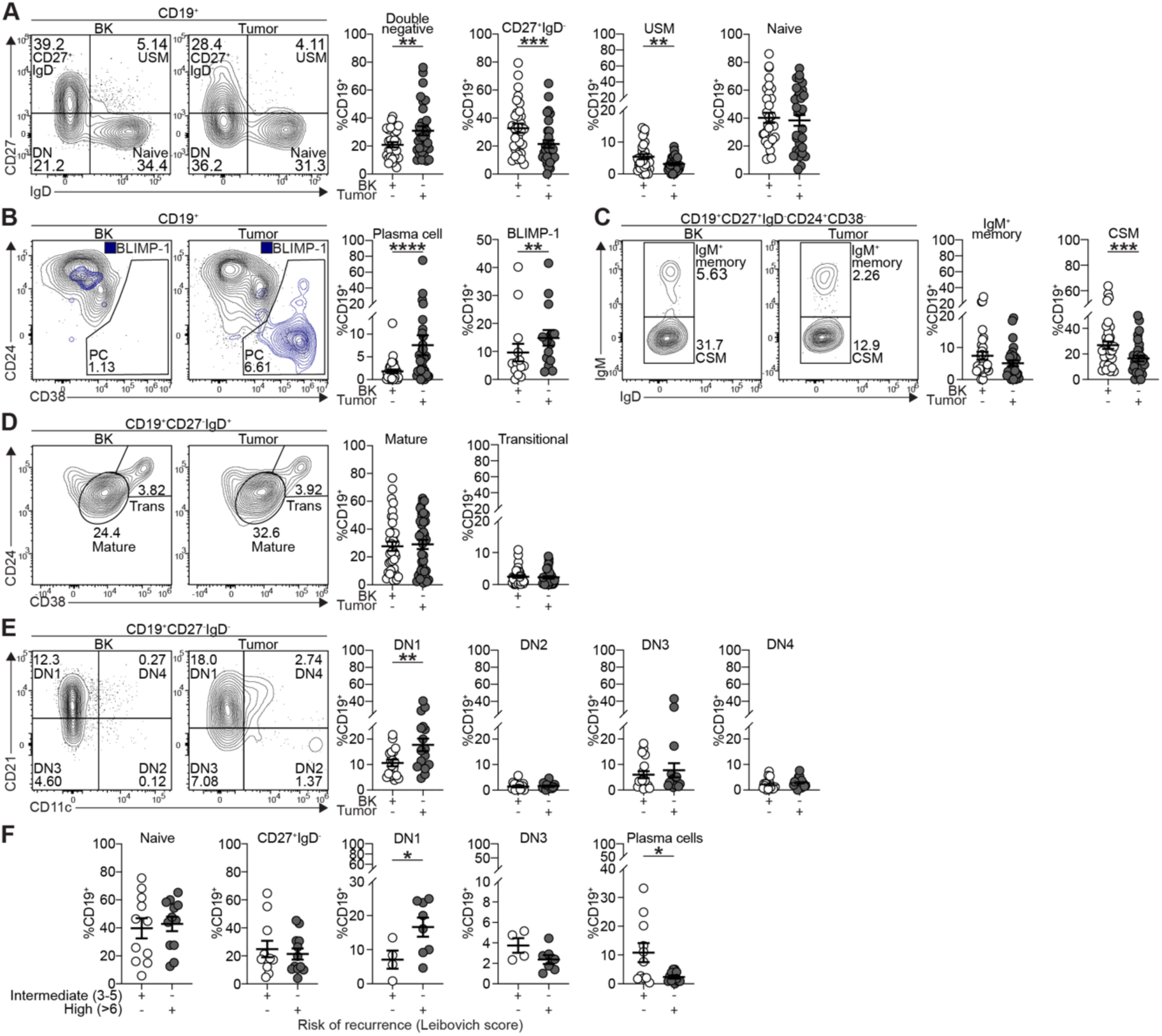
Increased DN1 B cell abundance in fresh tumor explants versus background tissue correlates with poor prognosis. (A) Representative contour plots and cumulative data showing the frequencies of CD19^+^CD27^-^IgD^-^ (double negative (DN)), CD19^+^CD27^+^IgD^-^, CD19^+^CD27^+^IgD^+^ (unswitched memory (USM)) and CD19^+^CD27^-^IgD^+^ (naive) B cells *ex vivo* in paired BK and matching tumor tissue; \*\**p*<0.01, \*\*\**p*<0.001. *n=*32 biologically independent samples. DN, USM, and naive B cells analyzed by paired two-tailed Student’s *t*-test, CD19^+^CD27^+^IgD^-^ B cells analyzed by paired two-tailed Wilcoxon test. Error bars represented as mean±SEM. (B) Representative contour plots and cumulative data showing the frequencies of CD19^+^CD24^lo/-^CD38^hi^ (plasma cells (PCs)) and BLIMP-1^+^ B cells *ex vivo* in paired BK and matching tumor tissue. BLIMP-1 expression overlaid onto the B cell contour plot; \*\**p*<0.01, \*\*\*\**p*<0.0001. For plasma cells *n*=31 and for BLIMP-1^+^ B cells *n*=15 biologically independent samples. Paired two-tailed Wilcoxon test. Error bars represented as mean±SEM. (C) Representative contour plots and cumulative data showing the frequencies of CD19^+^CD27^+^IgD^-^CD24^+^CD38^-^IgM^+^ (IgM^+^ memory) and CD19^+^CD27^+^IgD^-^CD24^+^CD38^-^IgM^-^ (class-switched memory (CSM)) B cells *ex vivo* in paired BK and tumor; \*\*\**p*<0.001. *n=*32 biologically independent samples. CSM analyzed by paired two-tailed Student’s *t*-test, IgM^+^ memory analyzed by paired two-tailed Wilcoxon test. Error bars represented as mean±SEM. (D) Representative contour plots and cumulative data showing the frequencies of CD19^+^CD27^-^IgD^+^CD24^int^CD38^int^ (mature) and CD19^+^CD27^-^IgD^+^CD24^hi^CD38^hi^ transitional B cells *ex vivo* in paired BK and tumor. *n=*32 biologically independent samples. Paired two-tailed Wilcoxon test. Error bars represented as mean±SEM. (E) Representative contour plots and cumulative data showing the frequencies of CD19^+^CD27^-^IgD^-^CD21^+^CD11c^-^ (DN1), CD19^+^CD27^-^IgD^-^CD21^-^CD11c^+^ (DN2), CD19^+^CD27^-^IgD^-^CD21^-^CD11c^-^ (DN3), and CD19^+^ CD27^-^IgD^-^CD21^+^CD11c^+^ (DN4) B cells *ex vivo* in BK and tumor; \*\**p*<0.01. *n=*18 biologically independent samples. Paired two-tailed Student’s *t*-test. Error bars represented as mean±SEM. (F) Cumulative data showing the frequencies of CD19^+^CD27^-^IgD^+^ (naive), CD19^+^CD27^+^IgD^-^, CD19^+^CD27^-^IgD^-^CD21^+^CD11c^-^ (DN1), CD19^+^CD27^-^IgD^-^CD21^-^CD11c^-^ (DN3) B cells, and CD19^+^CD24^lo/-^CD38^hi^ (plasma cells (PCs)) stratified according to intermediate (3-5) or high (>6) Leibovich scores; \**p*<0.05, \*\**p*<0.01. For naive, CD27^+^IgD^-^, and plasma cells *n=*26, for DN1 and DN3 *n=*13 biologically independent samples. Naive, DN1, DN3, and PC analyzed by unpaired two-tailed Student’s *t*-test, CD27^+^IgD^-^ analyzed by unpaired Mann-Whitney U test. Error bars represented as mean±SEM.

Using the validated Leibovich score to predict recurrence risk following nephrectomy^31^, we show that high-risk individuals have higher levels of DN1 B cells, while intermediate-risk individuals have more DN3 B cells and plasma cells (Figure 3F). These data suggest that DN1 B cells could serve as a prognostic measure of tumor recurrence in RCC.

### DN1 B cells are located within immature TLSs in RCC

TLS maturation status has been previously recognized as a strong predictor of clinical outcome, with mature TLSs associated with improved survival and better responses to immunotherapy^32,33^. In contrast, immature TLSs, linked to immunosuppression and T cell exhaustion, are associated with poorer survival and increased tumor progression^34^. Hematoxylin and Eosin (H&E) staining imaging of tumor sections from a retrospective cohort of 3 treatment-naive pre-metastatic RCC patients (Table S4B) shows a significant higher number of lymphoid aggregates in the non-cancerous supportive tissue defined as the tumor stroma compared to the tumor core, perinephric fat, or BK, and are more frequent in central sinus fat than perinephric regions (Figure 4A and 4B). Consistent with this, our RCC scRNA-seq dataset shows that the 12-chemokine TLS signature, a validated predictor of TLS presence^35^, is enriched in central tumor tissue relative to the tumor margin, BK, and blood (Figure 4C). High-dimensional spatial fluorescence analysis of selected lymphocyte-enriched regions of interest (ROIs) reveal B and T cell enriched areas, adjacent to PNAd⁺ vasculature, confirming TLS identity (Figure 4D). However, the lack of organized network of CD21^+^CD35^+^follicular dendritic cells (FDC), scarcity of CD4^+^PD-1^+^CXCR5^+^T follicular helper (TFH) cells, and the virtual absence of CD20^+^K*i*-67⁺ proliferating B cells is more representative of immature TLSs (Figure 4D and 4E). Plasma cells are located outside the immature TLSs, interspersed within tumor and adipose tissues (Figure 4D).

**Figure 4.**
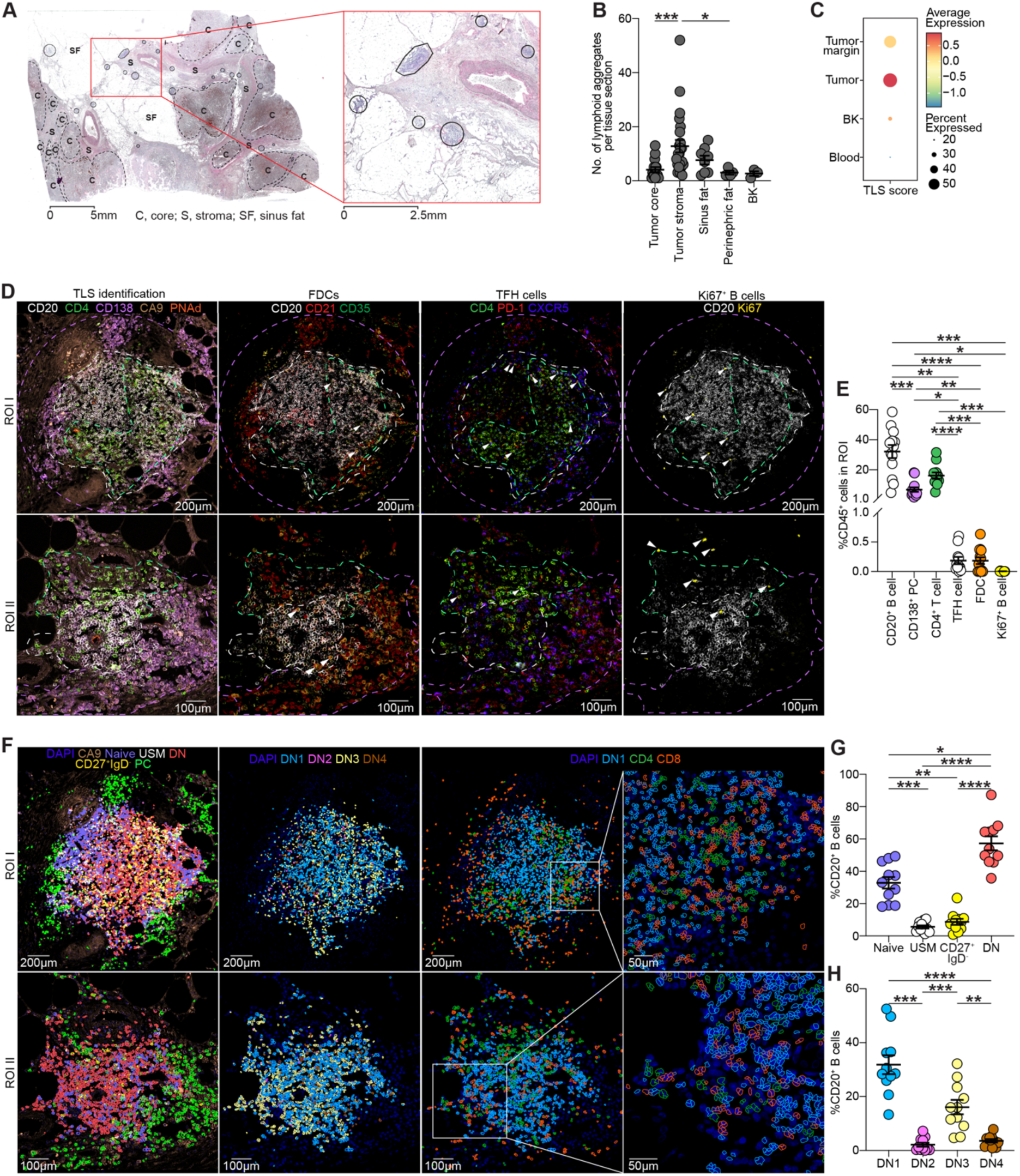
DN1 B cells are enriched within immature TLSs. (A) A representative image of H&E-stained tumor showing lymphoid aggregates in the in the core (C), stroma (S), sinus fat (SF). Scale bar = 5mm for whole section (left) and 2.5mm for the enlarged region (right). (B) Graph showing the number of lymphoid aggregates identified within the RCC tumor core and surrounding tissues; \**p*<0.05, \*\*\**p*<0.001. 38 tumor sections and 3 paired BK sections across *n*=3 biologically independent samples. Kruskal-Wallis test with Dunn’s test for multiple comparisons. Error bars represented as mean±SEM. (C) Dot plot showing the proportion of cells expressing a published 12-chemokine TLS signature^35^ (*CCL2, CCL3, CCL4, CCL5, CCL8, CCL18, CCL19, CCL21, CXCL9, CXCL10, CXCL11*, and *CXCL13*) (dot size), and their average expression levels (color intensity) in the RCC tumor, tumor margin, BK, and blood. (D) Spatial immunofluorescent image showing two representative ROIs (ROI I and ROI II) of tertiary lymphoid structures (TLSs) stained with CD20 (white), CD4 (green), CD138 (purple), CA9 (brown), and PNAd (orange) for TLS identification (Panel 1); CD20 (white), CD21 (red), and CD35 (green) for follicular dendritic cell (FDC) identification (Panel 2); CD4 (green), PD-1 (red), and CXCR5 (blue) for T follicular helper (TFH) cell identification (Panel 3); and CD20 (white) and K*i-*67 (yellow) for proliferating GC B cell identification (Panel 4). Scale bar = 200µm (ROI I), 100µm for (ROI II). (E) Cumulative data showing the frequencies of CD20^+^ B cells, CD138^+^ plasma cells, CD4^+^ T cells, CD4^+^PD-1^+^CXCR5^+^ TFH cells, CD21^+^CD35^+^ FDCs, and CD20^+^K*i*-67^+^ B cells within ROIs; \**p*<0.05, \*\**p*<0.01, \*\*\**p*<0.001, \*\*\*\**p*<0.0001. 13 ROIs across *n*=3 biologically independent samples. Kruskal-Wallis test with Dunn’s test for multiple comparisons. Error bars represented as mean±SEM. (F) Spatial analysis of B cells subsets in the TLS for two representative ROIs (ROI I and ROI II). Staining of DAPI (dark blue) and CA9 (brown) overlaid with cell segmentation (gating strategy shown in Figure S5A) of CD20^+^CD27^-^IgD^+^ (naive) (purple), CD20^+^CD27^+^IgD^+^ (unswitched memory (USM)) (white), CD20^+^CD27^-^IgD^-^ (double negative (DN)) (pink), CD20^+^CD27^+^IgD^-^ B cells (yellow), and CD138^+^ plasma cells (green) (Panel 1). DAPI staining (dark blue) plus cell segmentation of CD20^+^CD27^-^IgD^-^CD21^+^CD11c^-^ (DN1) (blue), CD20^+^CD27^-^IgD^-^CD21^-^CD11c^+^ (DN2) (pink), CD20^+^CD27^-^IgD^-^CD21^-^CD11c^-^ (DN3) (yellow), and CD20^+^CD27^-^IgD^-^CD21^+^CD11c^+^ (DN4) (brown) (Panel 2). DAPI staining (dark blue) plus cell segmentation of CD20^+^CD27^-^IgD^-^CD21^+^CD11c^-^ (DN1) (blue), CD4^+^ T cells (green), and CD8^+^ T cells (orange) (Panels 3 and 4). Panels 1-3 scale bar = 200µm (ROI I), 100µm for (ROI II), Panel 4 scale bar = 50µm. (G) Cumulative data showing the frequencies of CD20^+^CD27^-^IgD^+^ (naive), CD20^+^CD27^+^IgD^+^ (unswitched memory (USM)), CD20^+^CD27^+^IgD^-^, and CD20^+^CD27^-^ IgD^-^ (double negative (DN)) B cells within ROIs; \**p*<0.05, \*\**p*<0.01, \*\*\**p*<0.001, \*\*\*\**p*<0.0001. 11 ROIs across *n*=3 biologically independent samples. One-way ANOVA with Tukey’s test for multiple comparisons. Error bars represented as mean±SEM. (H) Cumulative data showing the frequencies of CD20^+^CD27^-^IgD^-^CD21^+^CD11c^-^ (DN1), CD20^+^CD27^-^IgD^-^CD21^-^CD11c^+^ (DN2), CD20^+^CD27^-^IgD^-^CD21^-^CD11c^-^ (DN3), CD20^+^CD27^-^IgD^-^CD21^+^CD11c^+^ (DN4) B cells within ROIs; \*\**p*<0.01, \*\*\**p*<0.001, \*\*\*\**p*<0.0001. 11 ROIs across *n*=3 biologically independent samples. One-way ANOVA with Tukey’s test for multiple comparisons. Error bars represented as mean±SEM.

Next, to address B cell subset location, we implemented cell segmentation (gating strategy in Figure S5A). Within TLSs, DN B cells are the most abundant population, followed by naive B cells, with CD27^+^IgD^-^ and unswitched memory (USM) subsets scarcely present (Figure 4F and 4G) confirming the scRNA-seq and flow cytometry results (Figure 2D and 3A). Among DN B cell subsets, DN1 are the most prevalent, followed by DN3 B cells, in the TLSs including in the T cell zone (Figure 4F and 4H).

### Bregs are enriched within immature TLSs

Both worse disease prognosis and TLS immaturity have been associated previously to increased number of regulatory cells, albeit most of the data are accrued from peripheral blood of cancer patients and not from intratumoral B cells^36–41^. Since B cells within TLS are known to participate more actively in immune modulation than their counterparts in the stroma or tumor core^42,43^, we focussed our investigation on these TLS-resident B cells.

Amongst tumor-resident B cells, DN1 B cells stand out for their distinctive enrichment of immunoregulatory genes associated with Breg function, including *IL10*, *TGFB1*, *TXN*, *AHR, HAVCR2, TIGIT, CD274*, and *PDCD1* (Figure 5A)^44,45^. A Breg module score, devised by averaging the expression of the Breg-associated genes, is also highly expressed in tumor, moderately in tumor margin, but negligibly expressed in peripheral B cells and in those residing in the BK (Figure 5B).

**Figure 5.**
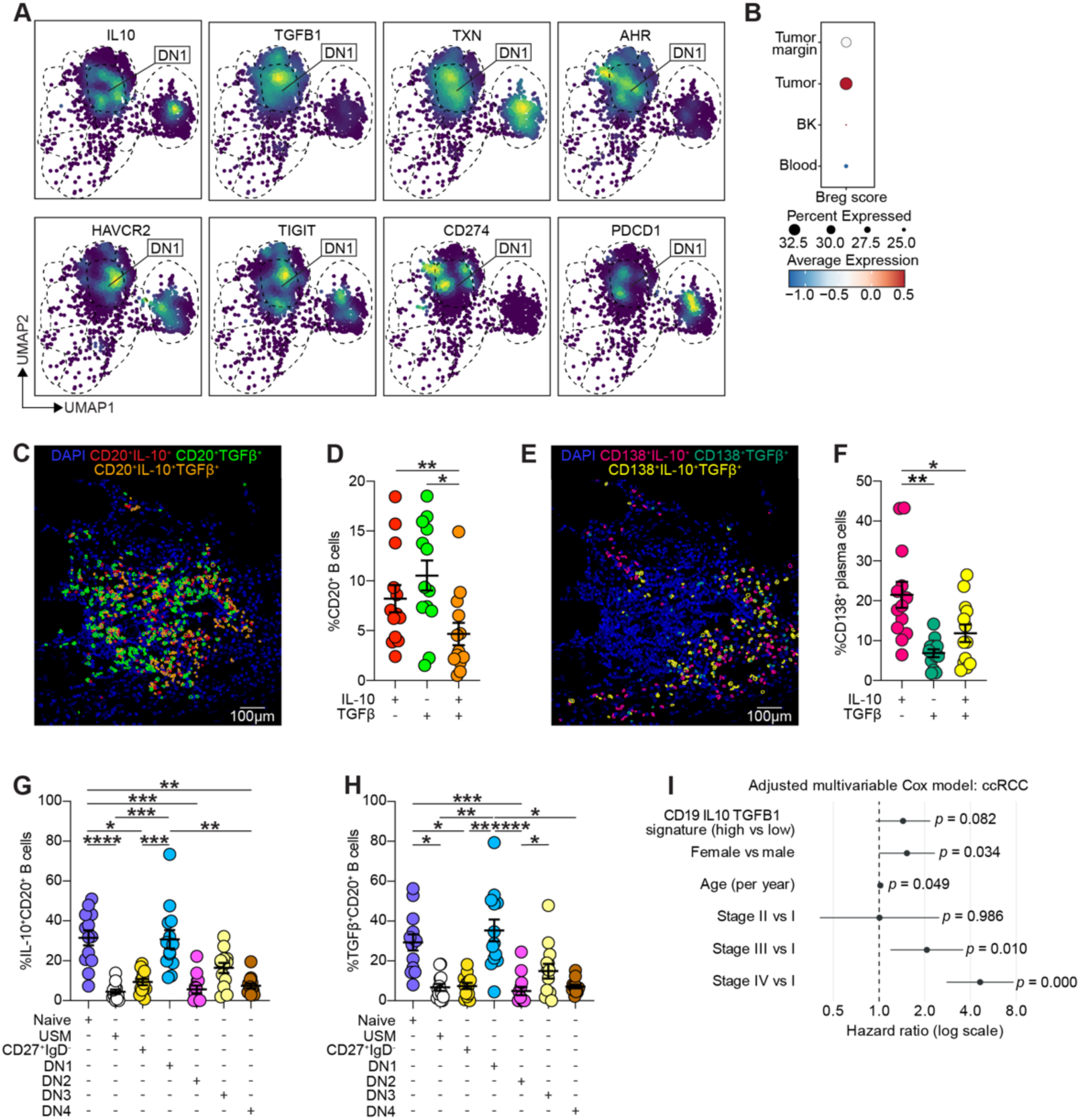
Bregs are enriched among naive and DN1 B cells in RCC and localize within immature TLSs. (A) Density UMAPs showing the distribution of *IL10^hi^, TGFB^hi^, TXN^hi^, AHR^hi^*, *HAVCR2^hi^, TIGIT^hi^*, *CD274^hi^ and PDCD1^hi^*RCC-resident B cells. (B) Dotplot showing the proportion of cells expressing a Breg score comprising *IL10, CD274, AHR, TXN, LAG3, TIGIT, CTLA4, CD86, TGFB1, and GZMB* (dot size), and their average expression levels (color intensity) in the tumor, tumor margin, BK, and blood. (C) Spatial analysis of segmented CD20^+^ Bregs in a representative TLS. Staining of DAPI (dark blue) overlaid with cell segmentation (gating strategy shown in Figure S5A) of CD20^+^IL-10^+^ Bregs (red), CD20^+^TGFβ^+^ Bregs (green), and CD20^+^IL-10^+^TGFβ^+^ Bregs (orange). Scale bar = 100µm. (D) Cumulative data showing the frequencies of CD20^+^IL-10^+^ Bregs, CD20^+^TGFβ^+^ Bregs, CD20^+^IL-10^+^TGFβ^+^ within ROIs; \**p*<0.05, \*\**p*<0.01. 13 ROIs across *n*=3 biologically independent samples. One-way ANOVA with Tukey’s test for multiple comparisons. Error bars represented as mean±SEM. (E) Spatial analysis of segmented CD138^+^ Bregs in a representative TLS. Staining of DAPI (dark blue) overlaid with cell segmentation (gating strategy shown in Figure S5A) of CD138^+^IL-10^+^ Bregs (pink), CD138^+^TGFβ^+^ Bregs (dark green), and CD138^+^IL-10^+^TGFβ^+^ Bregs (yellow) within TLSs. (F) Cumulative data showing the frequencies of CD138^+^IL-10^+^ Bregs, CD20^+^ CD138^+^TGFβ^+^ Bregs, CD138^+^IL-10^+^TGFβ^+^ Bregs within ROIs; \**p*<0.05, \*\**p*<0.01. 13 ROIs across *n*=3 biologically independent samples. One-way ANOVA with Tukey’s test for multiple comparisons. Error bars represented as mean±SEM. (G) Cumulative data showing the frequencies of CD20^+^CD27^-^IgD^+^ (naive), CD20^+^CD27^+^IgD^+^ (unswitched memory (USM)), CD20^+^CD27^+^IgD^-^ B cells, CD20^+^CD27^-^IgD^-^CD21^+^CD11c^-^ (DN1), CD20^+^CD27^-^IgD^-^CD21^-^CD11c^+^ (DN2), CD20^+^CD27^-^IgD^-^CD21^-^CD11c^-^ (DN3), and CD20^+^CD27^-^IgD^-^CD21^+^CD11c^+^ (DN4) within CD20^+^IL-10^+^ Bregs; \**p*<0.05, \*\**p*<0.01, \*\*\**p*<0.001, \*\*\*\**p*<0.0001. 13 ROIs across *n*=3 biologically independent samples. Fridman test with Dunn’s test for multiple comparisons. Error bars represented as mean±SEM. (H) Cumulative data showing the frequencies of CD20^+^CD27^-^IgD^+^ (naive), CD20^+^CD27^+^IgD^+^ (unswitched memory (USM)), CD20^+^CD27^-^IgD^-^ (double negative (DN)), CD20^+^CD27^+^IgD^-^ B cells, CD20^+^CD27^-^IgD^-^CD21^+^CD11c^-^ (DN1), CD20^+^CD27^-^ IgD^-^CD21^-^CD11c^+^ (DN2), CD20^+^CD27^-^IgD^-^CD21^-^CD11c^-^ (DN3), and CD20^+^CD27^-^IgD^-^CD21^+^CD11c^+^ (DN4) B cells within CD20^+^TGFβ^+^ Bregs; \**p*<0.05, \*\**p*<0.01. 13 ROIs across *n*=3 biologically independent samples. Fridman test with Dunn’s test for multiple comparisons. Error bars represented as mean±SEM. (I) Prognostic association of high versus low *CD19, IL-10, TGFB1* signature with survival in RCC using The Cancer Genome Atlas (TCGA) data. *p* values were derived by Cox proportional hazard model adjusting for signature group, age, sex, and American Joint Committee on Cancer (AJCC) stage.

Given the immunoregulatory signature characteristic of DN1 B cells, their localization within TLS, and their association with poorer prognosis, next we hypothesize that DN1 B cells play a role in suppressing anti-tumor immune responses. Spatial cell segmentation confirms that CD20⁺B cells express IL-10, TGFβ, or both, within immature TLSs (Figure 5C and 5D), whereas CD138⁺plasma cells expressing primarily IL-10 are also confined to the periphery of TLS (Figure 5E and 5F). Phenotypic analysis of TLS-residing cytokine-producing IL-10^+^ and TGFβ^+^Bregs are predominantly found within the naive and DN1 B cell compartments (Figure 5G and 5H). To assess their clinical relevance, we then defined a *CD19, IL10 and TGFB1* signature using TCGA data and applied an adjusted multivariable Cox proportional hazards model. High signature scores were associated with reduced overall survival (hazard ratio (HR) 1.44, 95% confidence interval (CI) 0.95-2.18) (Figure 5I).

### TLR signaling drives DN1 Breg differentiation

To investigate whether tumor-derived “cues” can drive DN1 B cell differentiation, we cultured healthy peripheral blood-derived naive B cells with several RCC cell lines. RCC lines induced a marked increase in DN B cells, whereas the HEK-293 control cell line, derived from an embryonic human kidney, promoted the expansion of memory B cell subsets (Figure S6). These findings suggest that RCC cells, either directly by cell contact or indirectly by releasing soluble factors, promote DN B cell differentiation.

Next, we sought to identify the specific extracellular signals released by RCC cells that may influence the emergence of DN1 Bregs. Unsurprisingly, gene set enrichment analysis (GSEA) identifies an upregulation of nucleic acid sensing Toll-like receptors (TLRs) as a stimulator for DN1 B cell clusters (Figure 6A), consistent with the established role of TLR family ligands in driving both DN B cell and Breg differentiation^9,29^. Notably, *TLR7* and *TLR9*, along with the downstream signaling mediators *MYD88, IRAK1/4, TRAF6, MAP3K7*, and *NFKB1*, were enriched within DN1 B cells (Figure 6B). The relevance of TLR7 and TLR9 in DN1 B cells expansion was also confirmed by flow cytometry (Figure S7A and S7B).

**Figure 6.**
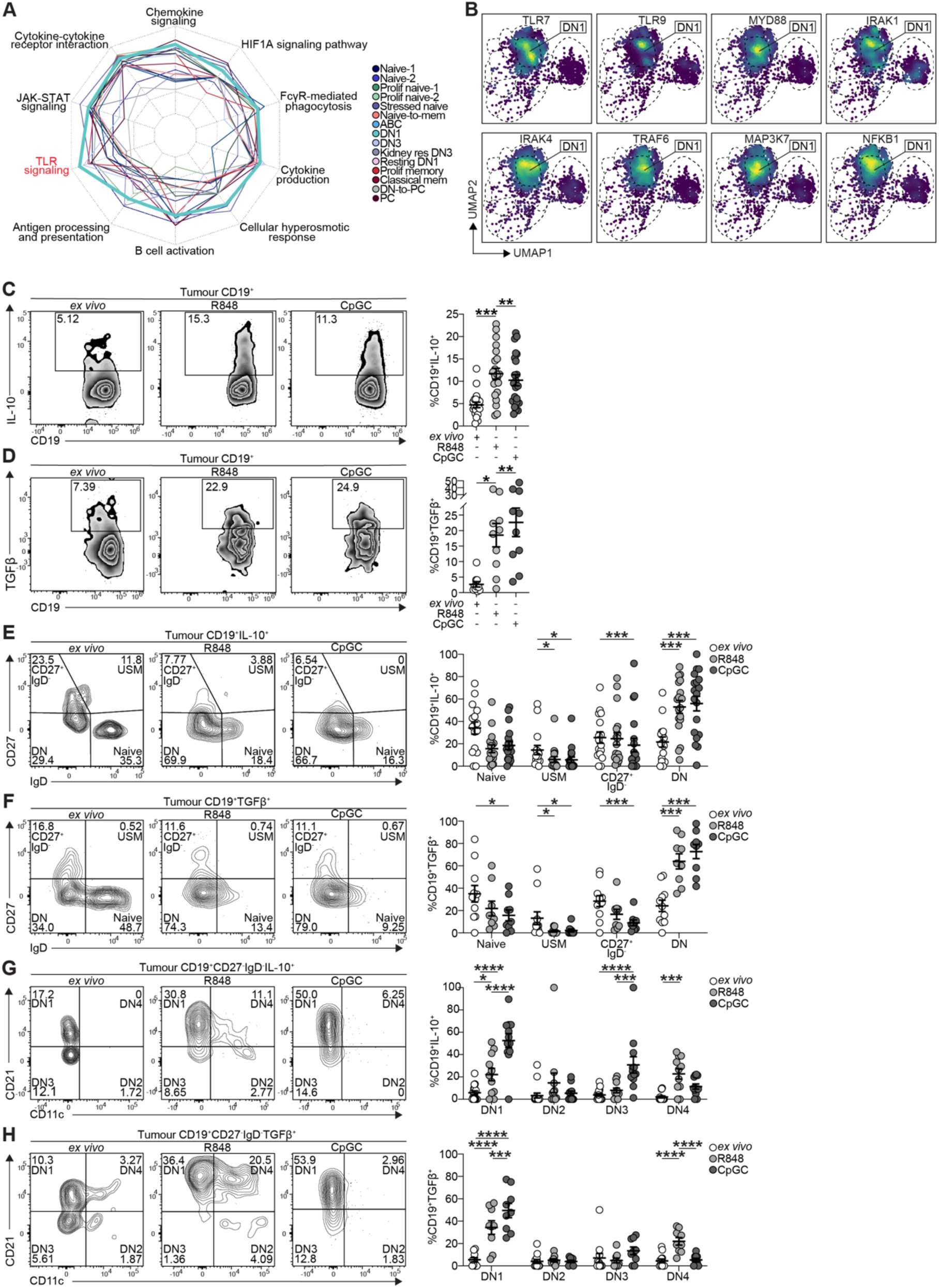
TLR7 and TLR9 drive DN1 Breg expansion. (A) Radar plot showing the Human Molecular Signatures Database and Kyoto Encyclopedia of Genes and Genomes (KEGG) pathways enriched in each tumor-infiltrating B cell (TIB) cluster. (B) Density UMAPs showing the distribution of *TLR7^hi^, TLR9^hi^, MYD88^hi^, IRAK1^hi^, IRAK4^hi^, TRAF6^hi^, MAP3K7^hi^*, and *NFKB1^hi^* RCC-resident B cells. (C) Representative contour plots and cumulative data showing IL-10 expression by tumor-infiltrating CD19^+^B cells *ex vivo* (white) and following 72h stimulation with R848 (light grey) or CpGC (dark grey); \*\**p*<0.01, \*\*\**p*<0.001. For *ex vivo n*=16, R848 and CpGC *n*=22 biologically independent samples. Mixed-effects one-way ANOVA with Dunnett’s test for multiple comparisons. Error bars represented as mean±SEM. (D) Representative contour plots and cumulative data showing TGFβ expression by tumor-infiltrating CD19^+^B cells *ex vivo* (white) and following 72h stimulation with R848 (light grey) or CpGC (dark grey); \**p*<0.05, \*\**p*<0.01. For *ex vivo n*=13, R848 and CpGC *n*=10 biologically independent samples. One-way Fridman test with Dunn’s test for multiple comparisons. Error bars represented as mean±SEM. (E) Representative contour plots and cumulative data showing the frequencies of CD19^+^CD27^-^IgD^+^ (naive), CD19^+^CD27^+^IgD^+^ (unswitched memory (USM)), CD19^+^CD27^+^IgD^-^, and CD19^+^CD27^-^IgD^-^ (double negative (DN)) B cells within tumor-infiltrating CD19^+^IL-10^+^B cells, *ex vivo* and following 72h stimulation with R848 or CpGC; \**p*<0.05, \*\**p*<0.01, \*\*\*\**p*<0.0001. *n*=24 biologically independent samples. Two-way ANOVA with Tukey’s test for multiple comparisons. Error bars represented as mean±SEM. (F) Representative contour plots and cumulative data showing the frequencies of CD19^+^CD27^-^IgD^-^CD21^+^CD11c^-^ (DN1), CD19^+^CD27^-^IgD^-^CD21^-^CD11c^+^ (DN2), CD19^+^CD27^-^IgD^-^CD21^-^CD11c^-^ (DN3) and CD19^+^ CD27^-^IgD^-^CD21^+^CD11c^+^ (DN4) B cells within tumor-infiltrating CD19^+^IL-10^+^B cells, *ex vivo* and following 72h stimulation with R848 or CpGC; \**p*<0.05, \*\*\**p*<0.001, \*\*\*\**p*<0.0001. *n*=18 biologically independent samples. Two-way ANOVA with Tukey’s test for multiple comparisons. Error bars represented as mean±SEM. (G) Representative contour plots and cumulative data showing the frequencies of CD19^+^CD27^-^IgD^+^ (naive), CD19^+^CD27^+^IgD^+^ (unswitched memory (USM)), CD19^+^CD27^+^IgD^-^, and CD19^+^CD27^-^IgD^-^ (double negative (DN)) B cells within tumor-infiltrating CD19^+^TGFβ^+^B cells, *ex vivo* and following 72h stimulation with R848 or CpGC; \*\**p*<0.01, \*\*\*\**p*<0.0001. *n*=12 biologically independent samples. Two-way ANOVA with Tukey’s test for multiple comparisons. Error bars represented as mean±SEM. (H) Representative contour plots and cumulative data showing the frequencies of CD19^+^CD27^-^IgD^-^CD21^+^CD11c^-^ (DN1), CD19^+^CD27^-^IgD^-^CD21^-^CD11c^+^ (DN2), CD19^+^CD27^-^IgD^-^CD21^-^CD11c^-^ (DN3) and CD19^+^ CD27^-^IgD^-^CD21^+^CD11c^+^ (DN4) B cells within tumor-infiltrating CD19^+^TGFβ^+^B cells, *ex vivo* and following 72h stimulation with R848 or CpGC; \*\*\**p*<0.001, \*\*\*\**p*<0.0001. *n*=11 biologically independent samples. Two-way ANOVA with Tukey’s test for multiple comparisons. Error bars represented as mean±SEM.

In parallel, gene expression profiling of tumor cells from our RCC scRNA-seq dataset shows elevated levels of necrosis-associated genes, including *HMGB1* and *ANXA5*, and reduced expression of apoptosis-related genes such as *CASP9* and *BCL2* (Figure S7C). This pattern is often observed under metabolic stress and hypoxic conditions within the TME^46–49^, corroborating the hypothesis that cell-free nucleic acids are released during necrotic tumor cell death to readily stimulate Bregs.

Given the established role of TLR stimulation in Breg differentiation in human peripheral blood^9^, we cultured purified TILs with surrogate ligands for TLR7 (R848) and TLR9 (CpGC), and assessed Breg frequencies and phenotype. On average, approximately 5% of tumor-resident B cells spontaneously express IL-10 or TGFβ (Figure 6C and 6D). Following TLR7 or TLR9 stimulation, the frequencies of CD19^+^IL-10^+^ and CD19^+^TGFβ^+^Bregs increase significantly in the tumor compared to *ex vivo* (Figure 6C and 6D). No significant expansion of LAG-3⁺ or PD-1⁺B cells is observed in the tumor compared to *ex vivo*, while PD-L1^+^Bregs were increase only in response to TLR7 (Figure S7D). Alternative known Breg stimuli^9^, including anti-IgM/IgG/IgA (anti-BCR) or CD40 ligand (CD40L), failed to induce comparable CD19^+^IL-10^+^Breg differentiation (Figure S7E).

*Ex vivo*, CD19⁺IL-10⁺ and CD19⁺TGF-β⁺Bregs are predominantly found within naive, CD27^+^IgD^-^ (including PCs), and double-negative (DN) B cells, and to a lesser extent within unswitched memory (USM) (Figure 6E, 6F), confirming our spatial Breg phenotyping results. In response to TLRs, CD19⁺IL-10⁺ and CD19⁺TGF-β⁺Bregs are predominantly enriched in the DN B cell compartment (Figures 6E and 6F), more specifically in the DN1 and, to a lesser extent, in DN4 B cells following stimulation with TLR7 (Figures 6G and 6H). These findings identify the DN1 B cell as a prominent inducible population within the RCC TME, with a subset exhibiting regulatory functions.

### DN1 Bregs are spatially associated with IL-10⁺ and TGFβ⁺CD8⁺T cells within TLSs

To explore the mechanisms by which DN1 Bregs may modulate local immunity, we assessed intercellular communication networks within the TME. Cell-cell interaction analysis revealed that B cells engage most robustly with CD8⁺T cells, compared to tumor cells or other immune subsets (Figure 7A and 7B). CellChat analysis further indicated that CD8⁺T cells predominantly act as signal recipients, with comparatively limited outgoing signaling capacity, highlighting their susceptibility to immune modulation by neighboring cell types (Figure 7C).

**Figure 7.**
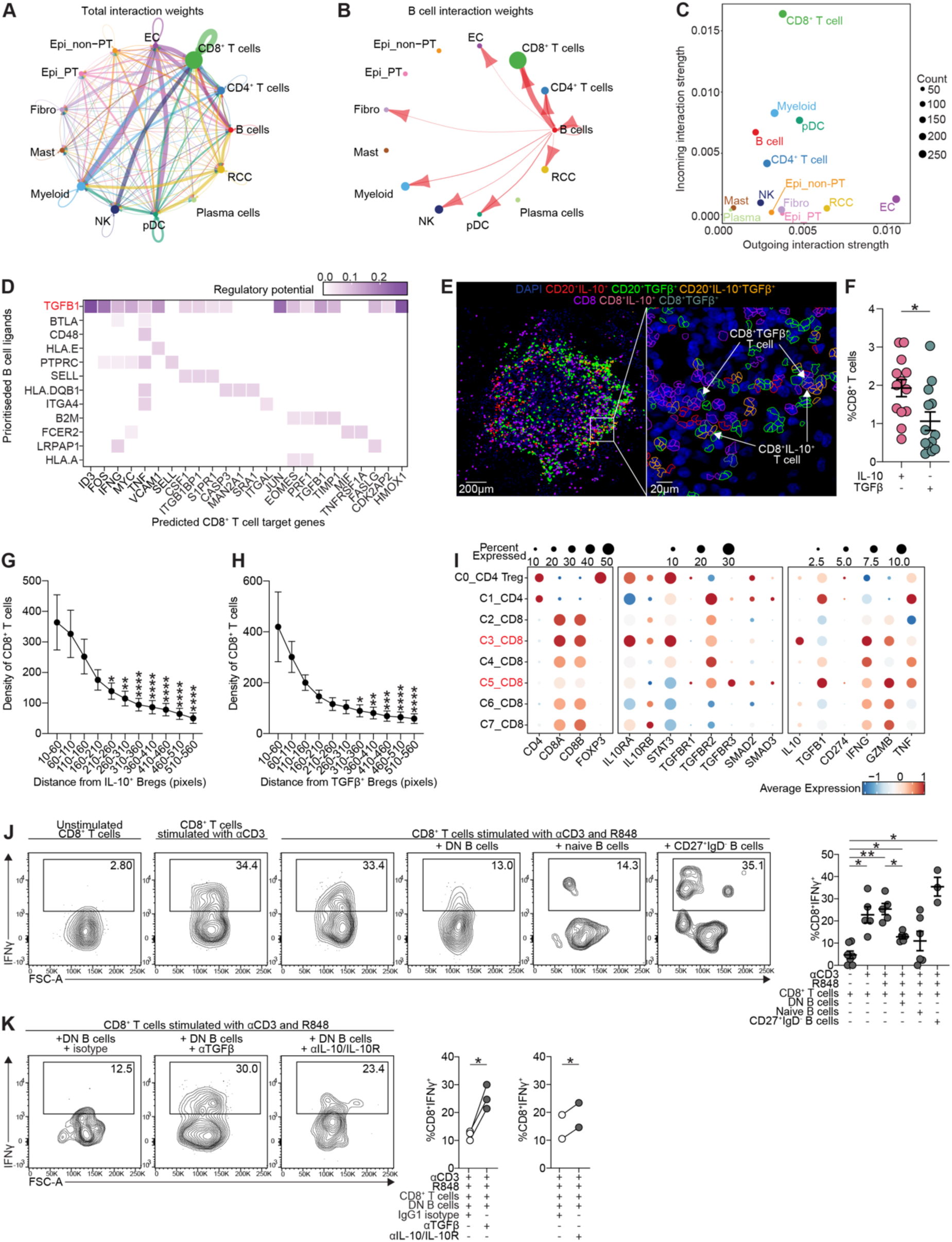
DN1 Bregs localize near IL-10⁺ and TGFβ⁺CD8⁺T cells, and DN B cells suppress IFNγ⁺CD8⁺T cell responses. (A) Circle plot of all RCC-resident cell types showing predicted outgoing ligand-receptor interactions, with edge width proportional to interaction strength. EC = endothelial cell; Epi_PT = proximal tubule epithelial cell; Epi_nonPT = non-proximal tubule epithelial cell. (B) Circle plot of RCC-resident B cells showing predicted outgoing ligand-receptor interactions, with edge width proportional to interaction strength. EC = endothelial cell; Epi_PT = proximal tubule epithelial cell; Epi_nonPT = non-proximal tubule epithelial cell. (C) Scatter plot showing the number of incoming versus outgoing interactions of RCC-resident cell types, calculated based on the aggregated communication probability across known ligand-receptor pairs. Dot size is proportional to the total interaction strength (incoming and outgoing) for each cell type. (D) Ligand-target heatmap showing the regulatory potential of the top 20 RCC-resident B cell ligands (rows) and their predicted target genes in RCC-resident CD8⁺ T cells (columns), with colour scale denoting regulatory potential. (E) Spatial analysis of segmented CD20^+^Bregs and CD8^+^T cells in a representative TLS. Staining of DAPI (dark blue) overlaid with cell segmentation (gating strategy shown in Figure S5A) of CD20^+^IL-10^+^ Bregs (red), CD20^+^TGFβ^+^ Bregs (green), CD20^+^IL-10^+^TGFβ^+^Bregs (orange), CD8^+^T cells (purple), CD8^+^IL-10^+^T cells (light pink) and CD8^+^TGFβ^+^T cells (grey). Scale bar = 200µm (Panel 1), 20µm for (Panel 2). (F) Cumulative data showing the frequencies of CD8^+^IL-10^+^T cells and CD8^+^TGFβ^+^T cells within TLSs; \**P*<0.05. 13 ROIs across *n*=3 biologically independent samples. Paired two-tailed Wilcoxon test. Error bars represented as mean±SEM. (G) The density of CD8^+^T cells within concentric bands of increasing distance in pixels from CD20^+^IL-10^+^ Bregs within TLSs; p<0.05, \*\**p*<0.01, \*\*\**p*<0.001, \*\*\*\**p*<0.0001. 13 ROIs across *n*=3 biologically independent samples. One-way Fridman test with Dunn’s test for multiple comparisons against the nearest band (10-60 pixels). Error bars represented as mean±SEM. (H) The density of CD8^+^T cells within concentric bands of increasing distance in pixels from CD20^+^TGFβ^+^Bregs within TLSs; \**p*<0.05, \*\**p*<0.01, \*\*\**p*<0.001, \*\*\*\**p*<0.0001. 13 ROIs across *n*=3 biologically independent samples. One-way Fridman test with Dunn’s test for multiple comparisons against the nearest band (10-60 pixels). Error bars represented as mean±SEM. (I) Dotplots showing the expression of indicated phenotypic and functional markers (dot size), and their average expression levels (color intensity) across RCC-resident T cell clusters. (J) Representative contour plots and cumulative data showing the frequencies of anti-CD3 and R848-stimulated IFNγ^+^CD8^+^T cells following a 72h co-culture with DN B cells, naive B cells, or CD27^-^IgD^+^B cells; \**p*<0.05, \*\**p*<0.01. *n=*7 biologically independent samples. Brown-Forsythe and Welch one-way ANOVA with Dunnett’s T3 test for multiple comparisons. Error bars represented as mean±SEM. (K) Representative contour plots and cumulative data showing the frequencies of anti-CD3 and R848-stimulated IFNγ^+^CD8^+^ T cells following a 72h co-culture with DN B cells in the presence of either an IgG1 isotype control, TGFβ-neutralizing antibody, or IL-10/IL-10-receptor (IL-10R) neutralizing antibodies; \**p*<0.05. For ɑTGFβ *n*=3 and ɑIL-10/IL-10R *n*=2 biologically independent samples. Paired two-tailed Student’s *t*-test.

Among the ligands expressed by B cells, TGFβ (*TGFB1*) emerges as the top regulatory molecule predicted to modulate the expression of cytotoxic effector genes (*IFNG*, *TNF*), transcriptional regulators (*EOMES*), and stress-response genes such as *HMOX1* in CD8⁺T cells (Figure 7D). These data suggest that DN1 Bregs may contribute to the functional reprogramming of CD8⁺T cells, potentially attenuating their cytotoxic activity. Spatial analysis shows that Bregs co-localize with CD8⁺T cells expressing IL-10 or TGFβ (Figure 7E and 7F). CD8⁺T cells are enriched within the concentric distance bands closest to CD20⁺IL-10⁺ and CD20⁺TGFβ⁺ Bregs across all ROIs analyzed, further supporting Breg-CD8^+^T cell proximity (Figure 7G and 7H). Unlike CD20⁺Bregs, CD138⁺IL-10⁺ and CD138⁺TGFβ⁺ plasma cell Bregs show limited co-localization with IL-10⁺ or TGFβ⁺CD8⁺ T cells (Figure S5B).

The close spatial association between IL-10^+^ and TGFβ⁺CD8^+^T cells and CD20^+^Bregs raises the possibility of a Breg-mediated conversion from effector to Treg phenotype, as previously reported^50^. We performed sub-clustering of CD4⁺ and CD8⁺T cells using our RCC scRNA-seq dataset, focusing on the expression of IL-10, TGFβ, their receptors, and associated downstream signaling components. Within the CD8⁺ compartment, cluster C3_CD8 co-expressed *IL10RA*, *IL10RB*, *STAT3*, and *IL10*, indicative of an IL-10 response loop (Figure 7I). Similarly, clusters C1_CD4 and C5_CD8 exhibited elevated expression of *TGFBR1*, *TGFBR2*, *TGFBR3*, *SMAD2*, and *SMAD3*, alongside *TGFB1*, consistent with active TGFβ signaling (Figure 7I). Collectively, these data support a model in which tumor-infiltrating DN1 B cells acquire regulatory functions in response to tumor signals, contributing to the formation of “immune-suppressive niches” within immature TLSs, and promoting the functional reprogramming of CD8⁺T cells, thereby facilitating immune evasion in RCC.

To examine whether tumor Bregs suppress IFNγ production by CD8⁺T cells, we depleted all B cells from both tumor and BK-derived single-cell suspensions. Cells were stimulated with anti-CD3 for 72 hours as previously shown^50^. In BK tissue, B cell depletion had no significant impact on the frequency of IFNγ⁺CD8⁺T cells (Figure S7F), whereas B cell depletion in tumor samples restores IFNγ expression in up to 40% of CD8⁺T cells, while TNFα and granzyme B (Gzmb) levels remain unchanged (Figure S7G and S7H).

To assess DN B cell suppressive capacity, we co-cultured anti-CD3 stimulated intra-tumoral CD8⁺T cells (to promote IFNγ production) with R848-stimulated purified autologous DN, naive, or CD27^+^IgD^-^ memory B cells for 72 hours. Co-culture with DN B cells, and to a lesser extent with naive B cells, which upon R848 stimulation differentiate into DN cells (Figure S7I), led to a reduction in IFNγ^+^CD8⁺T cells (Figure 7J). In contrast, CD27^+^IgD^-^B cells did not suppress IFNγ, confirming functional heterogeneity among B cell subsets (Figure 7J). Both TGFβ and IL-10 antibody neutralization partially rescues IFNγ expression in DN B cell co-cultures (Figure 7K), consistent with our *in silico* prediction that tumor-infiltrating B cells regulate CD8⁺T cell effector function via *TGFB1* (Figure 7D). Due to technical constraints, we were unable to directly isolate DN1 B cells in sufficient numbers. However, given that DN1 B cells account for the majority of DN subsets, it is plausible to speculate that they account for much of the observed regulatory activity. These findings reveal novel insights into B cell-mediated immunoregulation within the RCC TME and emphasizes the relevance of subset-specific analysis in understanding their contributions to shaping anti-tumor immunity.

## Discussion

B and plasma cells comprise 3%-30% of tumor-infiltrating immune cells, similar to T cells, although this proportion is highly variable, depending on cancer type, disease stage, and detection methodology^51^. The functional impact of B cells on tumor progression is cancer and therapy-dependent: they may exert either pro- or anti-tumorigenic effects based on their phenotypic state, antibody isotype, cytokine production, spatial distribution, and the nature of the tumor and its microenvironment^52^. In cancers like melanoma, lung adenocarcinoma, pancreatic adenocarcinoma, and head and neck squamous cell carcinoma (HNSCC), elevated B cell/plasma cell gene signatures correlate with improved survival^52^. In contrast, brain low-grade glioma and RCC show the opposite trend^52^, suggesting that B cell function is highly context-dependent and shaped by the tumor-specific immune milieu. The increase in number of pan-cancer B cell atlas studies, which merge TIBs across cancer types, have begun to fill in this knowledge gap by identifying commonalities in B cell subsets amongst different cancers^19–21^. However, integrated clustering may obscure subtle but physiologically relevant differences within one type of cancer, particularly in B cell-underrepresented tumor types. Our computational pipeline, which infers TIB states within individual cancer types and projects them into a shared latent space, highlights the pivotal influence of tissue- and cancer-specific microenvironments in shaping B cell phenotypic diversity and subset composition within tumors. By leveraging this framework, followed by validation in an independent scRNA-seq dataset, flow cytometry analysis, and spatial profiling, we identify a previously unrecognized enrichment of DN1 B cells in RCC, a cancer type typically characterized by low immune infiltration^20^.

DN B cells have been extensively characterized in systemic lupus erythematosus (SLE), and more recently in the context of COVID-19, where they have consistently been linked to pathogenic roles^29,53,54^. In cancer, however, the role of DN B cells remains less defined. IFN-induced B cells, including naive B cells, and DN B cells were enriched in nasopharyngeal carcinoma, although no prognostic information was given^55^. Similarly, DN B cells were found to be expanded in non-small cell lung cancer (NSCLC) and to negatively correlate with the number of memory B cells, which was associated with better tumor differentiation^56^. In HNSCC, EF-derived DN3 B cells are more prevalent in the TILs of stage T2 and T4 patients, suggesting a worse prognosis effect of these subsets^57^. In melanoma, blood-resident DN3 B cells were enriched in patients with worse survival^57^. ABCs have been the focus of considerable debate due to their contrasting associations with cancer prognosis. On one hand, ABCs have been linked to poorer survival outcomes in several solid tumors, including colon, stomach, lung, and liver cancers^21^. On the other hand, increased of FCRL4⁺ ABC B cells, are associated to improved survival across multiple cancer types, suggesting potential functional heterogeneity within the ABC population^20^. Our study adds a new layer of complexity to the evolving understanding of DN B cells in cancer. Clinically, only DN1 B cells, and no other B cell subset present in tumor, were correlated with a higher risk of recurrence in RCC, suggesting their potential utility as negative prognostic biomarkers and potentially opening new avenues for therapeutic targets. To our knowledge, this is the first report assigning a functional role to DN1 B cells, which have thus far been predominantly identified in the peripheral blood of healthy individuals^29^.

Using high-plex spatial immunofluorescence, we show that DN1 Bregs, enriched for immunosuppressive mediators (IL-10 and TGFβ) localize within immature TLSs nearby IL-10^+^ and TGFβ^+^CD8⁺T cells. Functionally, *in vitro* assays demonstrate that tumor derived DN B cells (highly enriched in DN1 B cells) suppress tumor IFNγ-producing CD8⁺T cell responses, in part through the secretion of IL-10 and TGFβ. Our findings, therefore, support previous reports hypothesizing that Bregs and Tregs co-localize in TLSs to dampen anti-tumor responses^38,58^. While the role of IL-10-expressing cells remains to be elucidated, their localization outside the TLS implies a spatially organized division of labour between CD20⁺ regulatory B cells and CD138⁺ plasma cell-like Bregs. Accumulating evidence from experimental cancer models has highlighted the pro-tumorigenic role of Bregs^59^. Our earlier work demonstrated that, during DMBA/TPA-induced squamous carcinogenesis, TNFα mediates tumor-promoting activity via Bregs, which suppress anti-tumor immune responses^60^. This finding has since been supported by several studies indicating that Bregs contribute to tumor development across various malignancies. Despite these findings in experimental model systems, robust evidence in humans remains limited, particularly regarding the intratumoral presence and immunosuppressive functions of Bregs. Notably, in breast cancer, CD19⁺IL-10⁺B cells have been detected within TIL aggregates and have been suggested to facilitate immunosuppression by inducing the conversion of resting CD4⁺T cells into regulatory T cells (Tregs)^10^. Similarly, in ovarian cancer, B cells have been identified in ascitic fluid, though their IL-10 production levels were very low, complicating interpretations of their regulatory potential^11^. Other studies have reported elevated levels of Bregs in the peripheral blood of patients with hepatocellular carcinoma^61^, breast cancer^62^, and esophageal squamous cell carcinoma^63^ compared to healthy controls. CD21^-^DN B cells have also been previously shown to correlate with a lack of response to checkpoint inhibitor (CI) therapy in NSCLC^64^. However, the mechanism by which CD21^-^DN B cells hamper CI therapy remains unknown, although it was hypothesized that, similarly to NSCLC, exhausted TIL-Bs (CD19^+^CD20^+^CD69^+^CD27^-^CD21^-^) could be associated with a Treg phenotype (FoxP3^+^CD4^+^)^2^. Therefore, to date, the role of Bregs intratumorally remains largely unknown.

The mechanisms driving Breg differentiation in tumors are not fully resolved. While Bregs can be induced by “homeostatic levels” of inflammatory stimuli, including TNFɑ^60^, IL-6 and IL1β^65^, tumors may promote Breg development through metabolites, exosomes, or necrotic debris^63,66,67^. Known inducers of Breg function, such as BCR engagement^68,69^, TLR ligation^67,68^, and CD40 signaling^70^, vary in relevance by context. In our study, DN1 B cells were enriched for TLR signaling pathways, and only TLR stimulation drove the differentiation of TIBs into the DN1 subset, with a proportion expressing immunoregulatory cytokines. This suggests that nucleic acid release from dying tumor cells may be a key driver of DN1 Breg differentiation in RCC, independently or co-aided by T cell help. This latter hypothesis is supported by the increase in the expression of genes predicting tumor necrosis, a process known to release nuclear acids; hypothesis that will be further investigated in future study.

In conclusion, while scRNA-seq has been instrumental in identifying previously unrecognized B cell subsets and refining our understanding of B cell populations in both tumors and adjacent tissues, computational predictions alone require experimental validation to determine functional relevance. This study presents a workflow that integrates scRNA-seq with spatial imaging and flow cytometry, enabling the identification of novel B cell populations and the elucidation of their roles in cancer-specific contexts. Our findings also reinforce the importance of cancer-type-specific B cell characterization, with significant implications for biomarker discovery and the development of targeted immunotherapies.

Limitations of the study

Although we employed computational strategies to minimize batch effects, technical variability inherent in scRNA-seq platforms and tissue processing as well as patient variability could influence subset detection and gene expression measurements.

Our study is based on cross-sectional single-cell RNA-seq data, which limits insights into the temporal dynamics and differentiation trajectories of B cell subsets, particularly under therapeutic pressure such as checkpoint inhibition.

While we observe distinct transcriptional profiles and prognostic associations, as for all human studies, the immunoregulatory roles of the CD27^-^IgD^-^CD21^-^CD11c^-^DN1 B cells in promoting pro-tumorigenic responses is limited to *in vitro* studies.

The association between DN1 B cells and poor response to checkpoint inhibitor therapy has not yet been established and warrants further investigation through prospective clinical validation in controlled immunotherapy trials.

While bulk TCGA survival data are often used to infer associations with gene signatures, the *CD27^lo^IGHD^lo^*phenotype of DN1 B cells made this approach less robust; accordingly, we did not include it in our analysis.

## Acknowledgments

This work is funded by: Biotechnology and Biological Sciences Research Council (BB/T002212/1) awarded to C.M. and F.F.; Biotechnology and Biological Sciences Research Council (BB/T008709/1) LIDo PhD studentship awarded to I.W., C.M., and F.F.; Medical Research Council DTP PhD studentship (MR/N031867/1) awarded to Z.B. and C.M.; GSK collaboration award awarded to C.M., M.G.B.T, and H.J.S.; Versus Arthritis UK program grant (21140) awarded to C.M. The Royal Free Hospital Biobank coordinated the collection of surgical specimens. Schematics were created with BioRender.com.

## Contributions

Z.B. and I.W. designed, performed experiments, analyzed data, and wrote the manuscript. C.J.M.P. and H.F.B. performed experiments and critically reviewed the manuscript. C.M. designed experiments, analyzed data, and wrote the manuscript. J.N. and F.F., designed experiments, analyzed data and critically reviewed the manuscript. T.J.M. provided scRNA-seq dataset and critically reviewed the manuscript. M.G.B.T. provided clinical samples, resources, and critically reviewed the manuscript. H.J.S. designed and critically reviewed the manuscript. A.R. provided clinical information and critically reviewed the manuscript. E.H.AB. provided clinical information. A.DT. critically reviewed data analysis and critically reviewed the manuscript.

## Competing interests

A.DT. is a full-time employee of GlaxoSmithKline and contributed to this work as part of a formal research collaboration funded by GlaxoSmithKline. A.DT. holds shares in the company.

## Experimental model and study participant details

### Human specimens

Paired RCC and BK samples were collected from radical nephrectomy surgeries at the Royal Free Hospital. Surgical specimens were either transported directly from the operating theatre to histopathology to minimize warm ischemia time or stored overnight in University of Wisconsin solution at 4°C before processing the following morning. Written informed consent was obtained for tissue and blood biobanking under the Characterization of the Immunological and Biological Markers of Renal Cancer Progression Study (NHS National Research Ethics Service reference 16/WS/0039). Pathology outcomes and clinical scores were collected retrospectively from electronic patient records, provided in Table S4A. Pre-cut slides of archived FFPE RCC and paired BK tissue sections from 3 patients were purchased from Health Services Laboratories, with clinical information provided in Table S4B. Blood cones were purchased from NHS Blood and Transplant under research ethics number 21/WA/0388.

### Cell lines

The human cell lines RCC4 and RCC10 were generously provided by Professor Maxine Tran, 786-O by Professor Marilena Loizidou, and HEK293T by Professor Hans Stauss. Cells were maintained in T-75 flasks in complete RPMI 1640 (Gibco) supplemented with 10% fetal bovine serum (FBS; LabTech) and 1% penicillin/streptomycin (P/S; Sigma-Aldrich), cultured in a humidified incubator at 37°C and 5% CO_2_. For twice-weekly passaging, culture media was aspirated, cells washed with PBS and incubated with 1X TrypLE Expression (Gibco) dissociation agent for 5 minutes. 90% of cells were discarded in each passage to avoid over-confluency. Mycoplasma testing of culture media was performed regularly.

## Method details

### Cell isolation and culture

Tumor and BK tissues were collected on ice in serum-free RPMI containing 1% P/S, washed in PBS containing CaCl_2_ and MgCl_2_ (Gibco), minced, and digested at 37°C for 15 min in 50μg/ml Liberase TL (Roche) in serum-free RPMI + 1% P/S with gentle agitation. Digests were filtered through 100μm and 70μm strainers, then lymphocytes were enriched using Percoll-based density centrifugation (40% isotonic Percoll underlaid with 80%).

Tissue-resident B cells were isolated using the EasySep immunomagnetic CD19 positive selection kit (STEMCELL, cat. no. 17854). The B cell-depleted filtrate was used for B cell depletion experiments. Peripheral blood mononuclear cells (PBMCs) were isolated from blood cones by Ficoll-based density centrifugation. B cells were isolated from PBMCs using the EasySep immunomagnetic negative selection kit (STEMCELL, cat. no. 19054). Tumor-infiltrating CD27^-^IgD^+^ naive, CD27^-^IgD^-^ DN, CD27^+^IgD^-^ B cells, and CD8^+^ T cells were purified by fluorescence-activated cell sorting (FACS) with a purity of >90%.

For immunophenotyping experiments, lymphocytes were stained *ex vivo* after isolation from tissue. For stimulation assays, lymphocytes or purified B cells were cultured in RPMI containing 10% FBS and 1% P/S at 37°C and 5% CO_2_ for 72h with either CpGC ODN 2395 (1μM, Invivogen), R848 (Resiquimod; 1μg/ml, Invivogen), mega-CD40L (1μg/ml, Enzo), or anti-IgM/IgG/IgA (10μ/ml, Sigma-Aldrich). To promote intracellular cytokine staining, cells were stimulated with phorbol 12-myristate 13-acetate (PMA; 50ng/ml, Sigma-Aldrich), ionomycin (250ng/ml, Sigma-Aldrich), and brefeldin A (5μg/ml, Sigma-Aldrich) for the final 5h of the culture. For assays involving the stimulation of T cells, 96-well U-bottom plates were pre-coated with 0.5μg/ml anti-CD3 (ThermoFisher Scientific) 2h at 37°C prior to plating cells. For suppression assays, purified tumor-infiltrating CD27^-^IgD^+^ naive, CD27^-^IgD^-^ DN, CD27^+^IgD^-^ B cells were cultured 1:1 with autologous CD8^+^ T cells for 72h with 1μg/ml R848 in culture plates pre-coated with 0.5μg/ml plate-bound anti-CD3.

### Flow cytometry

Single-cell suspensions (excluding the unstained control) were stained with LIVE/DEAD™ Fixable Blue Dead Cell Stain (ThermoFisher Scientific) at 1:500 and Human FcR Blocking Reagent (Miltenyi Biotec) at 1:10 in PBS for 20 minutes at room temperature. Surface markers were stained in Brilliant Stain Buffer (BD) for 20 minutes at 4 °C. For intracellular staining, cells were fixed and permeabilized using the intracellular fixation & permeabilization buffer kit for 20 minutes (ThermoFisher Scientific), and for intranuclear staining, cells were fixed and permeabilized using the FoxP3 Fixation buffer kit for 30 minutes (ThermoFisher Scientific). Intracellular and intranuclear antibodies were incubated for 40 minutes at 4 °C. Data were acquired on an Aurora spectral flow cytometer (Cytek) using spectral unmixing and subtraction of autofluorescence based on matched unstained controls. Analysis was performed using FlowJo (BD). Antibodies are detailed in Table S5.

### Hematoxylin and eosin (H&E) staining

H&E staining was performed using the Tissue-Tek DRS automated slide stainer (Sakura). Stained slides were scanned at 40X magnification using the NanoZoom S360 Digital Slide Scanner (Hamamatsu) and analyzed using NDP.view2 (Hamamatsu). Lymphoid aggregates were counted and averaged across two independent researchers.

### MACSima imaging cyclic staining (MICS)

Sections were deparaffinized in xylene, followed by rehydration through a series of graded ethanol dilutions (100%, 90%, 70%). Antigen retrieval was performed by immersing the slides in IHC-Tek Epitope Retrieval Solution (IHC World) and steaming for 40 minutes. A MACSwell^TM^ One imaging frame (Miltenyi Biotec) was mounted on the slide and tissue sections blocked with 5% normal donkey serum (Sigma-Aldrich) and permeabilized with 0.05% Triton-X (Sigma-Aldrich) for 20 minutes at room temperature. Sections were incubated with primary antibodies (Table S6) for 2 hours at room temperature, then pre-stained with DAPI (MACSima stain support kit, Miltenyi Biotec) at 1:5 in running buffer for 10 minutes at room temperature. Conjugated antibodies were prepared in a MACSWell^TM^ DeepWell Plate (Miltenyi Biotec) at appropriate dilutions in MACSima running buffer (Miltenyi Biotec) (Table S6) and loaded into the MACSima instrument (Miltenyi). A bleach cycle was first applied to minimize tissue autofluorescence. Sections were stained iteratively with FITC-, PE-, or APC-conjugated anti-human antibodies. After each cycle, photobleaching was performed to remove the fluorescence signal. Image acquisition and processing were performed by the MACSima instrument. Images were pre-processed using the MACS IQ View (Miltenyi).

Cell segmentation based on DAPI-stained nuclei was performed using the Advanced Tissue Morphology method with the confined donut algorithm. Cell populations were gated according to marker expression in MACS iQ View, with gate placement verified by overlaying segmented populations onto their corresponding immunofluorescent staining.

### scRNA-seq data assembly and pre-processing

ScRNA-seq datasets were obtained from the Ma *et al.* cancer B cell atlas (http://pancancer.cn/B/)^21^ and the Fitzsimons *et al.* cancer B cell atlas (https://doi.org/10.5281/zenodo.13385122)^19^. The atlases were merged into a single AnnData object and processed in scVI^71^ using the scanpy python package^72^ (version 1.10.1), removing any duplicated entries. B cells from BRCA, LC, COAD, and RCC were selected. Cells were annotated using the automated CellTypist annotation model^73^ and those identified as non-B cells, including plasma cells, were excluded. Patients with <70 cells and datasets with <400 cells were excluded to enhance the quality of integration. One RCC patient was excluded due to an abnormal BCR mutation rate distribution^21^. A mitochondrial threshold of 10% was used in line with the published Ma *et al.* dataset^21^. Each dataset was normalized to 10,000 counts per cell and log transformed. 8,000 highly variable genes were identified using the *sc.pp.highly_variable_genes* function (Seurat v3 method) implemented in scanpy, accounting for batch effects by specifying the dataset as the batch key.

Our in-house scRNA-seq RCC dataset (https://www.sanger.ac.uk/project/microenvironment-of-kidney-cancer/)^27^ was processed separately as above, using a 25% mitochondrial threshold to prevent loss of rare and stressed B cell populations.

### Data integration, dimension reduction, and unsupervised clustering

The cancer-specific scVI models and RCC-specific scVI model were trained using default parameters and a batch size of 130. The scVI batch-corrected gene expression values were generated by the *get_normalized_expression* function and scaled to a library size of 10,000. A k-nearest neighbors graph was computed using *sc.pp.neighbors*, specifying the scVI latent representations. UMAP embeddings were generated using *sc.tl.umap* to visualize the data in two dimensions, with the default distance parameter. Leiden clustering was applied using *sc.tl.leiden* with a default resolution parameter to generate the clusters.

The cancer-specific AnnData objects were concatenated into a single dataset and reprocessed following the same normalization, log transformation, and scaling steps. 10,000 highly variable genes were identified in this combined dataset, accounting for batch effects with a cancer_dataset batch key, an identifier that combines each patient’s cancer type and dataset ID. To optimize scVI’s latent-space inference across the integrated dataset of four cancer types, ribosomal protein genes were excluded from the highly variable gene set. A second scVI model was trained on the combined dataset to generate a shared latent space capturing intra- and inter-tumor heterogeneity. The scVI model was configured using the raw counts layer and included the cancer_dataset and patient as categorical covariates.

Cluster centroids were computed from scVI latent coordinates by averaging across cells in each B cell cluster. Pairwise cosine distances between centroids were calculated using *pdist* from SciPy, and hierarchical clustering was performed using the *linkage* function to generate a dendrogram of cluster similarity.

The pairwise cosine distances between cluster centroids helped to organize clusters into broad B cell groups. We then evaluated known B cell marker genes to validate these groupings and define their biological identity. Marker genes for each broad group were identified by comparing cells within that group, either against a lineage-matched reference or against all other B cell clusters, using a Wilcoxon rank-sum test. Genes were ranked by adjusted p-value and fold-change.

For pan-cancer clustering, we applied Leiden clustering with default parameters to the combined dataset’s scVI latent space, as described above, using a cut-off of 10 cells minimum per cluster. We then generated a contingency table showing the proportion of cells from each cancer-specific cluster that mapped to each pan-cancer cluster, quantifying overlap and assessing how faithfully the pan-cancer integrated clusters recapitulate the original cancer-specific groupings.

### BCR repertoire analysis

To compare somatic hypermutation frequencies across annotated B cell subsets and plasma cell populations, we integrated single-cell BCR sequencing (scBCR-seq) data from the Ma *et al.* atlas (http://pancancer.cn/B/). Mutational frequency values were aligned with our cross-cancer B cell atlas by merging the dataset with our annotated AnnData object. For comparison with plasma cells, the scBCR-seq dataset was also merged with the original Ma *et al.* Seurat object, which included pre-annotated plasma cells, and converted to AnnData for joint analysis.

### Comparison of gene expression to healthy PBMCs

To integrate the reference peripheral blood B cell dataset from Stewart *et al.*^26^ with the cancer-specific single-cell datasets (BRCA, LC, COAD, RCC), we first concatenated the individual AnnData objects for each cancer type with the blood B cell dataset into a single combined AnnData object. Normalization, logarithmic transformation, and scaling procedures were applied to the combined dataset, in line with the previously described workflows. A new scVI model was trained on the combined dataset to embed all cells in a shared latent space. We then extracted the denoised layer and assessed the median gene expression of *CD27* and *IGHD* across all clusters.

### Trajectory analysis

AnnData objects were converted into RDS files for Seurat^74^ (v5) using the sceasy^75^ package. Pseudotime analysis was performed using the Monocle 3 package^76^. Unsupervised clustering was applied using *cluster_cells()* prior to fitting a principal graph using *learn_graph()*. Based on prior knowledge, we chose naïve B cells as the root node of the trajectory. Cells were visualized and colored by “pseudotime” using *plot_cells()*.

### Pathway analysis

The fsgea package^77^ was used to perform pathway analysis. Upregulated pathways were identified among the top DEGs within each cluster. The Kyoto Encyclopaedia of Genes and Genomes (KEGG) dataset in the MSigDB package^78^ was used to identify significant pathways (*p*<0.05). The code for the radar plot was adapted from Fitzsimons *et al.*^19^.

### Cell-cell communication analysis

We used CellChat^79^ to investigate the number and strength of interactions between cell types present within RCC. Processed expression profiles from the data slot of the Seurat object, alongside cell type annotations, were used to create a CellChat object. The CellChatDB.human ligand-receptor interaction database was used to identify interactions.

NicheNet^80^ was used to further dissect gene regulatory interactions. B cells were selected as the sender cell type and CD8^+^ T cells as the receiver. NicheNet calculated regulatory potential scores for ligand-target pairs, identifying significant ligand-receptor-target interactions. The paired RCC blood dataset was used to identify background genes subtracted from significant receptor-ligand pairs observed in the tumor.

### The Cancer Genome Atlas (TCGA) data analysis

We obtained TCGA CIBERSORT^81^ immune fractions (“TCGA.Kallisto.fullIDs.cibersort.relative.tsv”) and leukocyte purity metrics (“TCGA_all_leuk_estimate.masked.20170107.tsv”) from the NCI Genomic Data Commons Data Portal. Naïve and memory B-cell fractions from CIBERSORT were summed and then multiplied by each sample’s ESTIMATE-derived leukocyte purity to yield the fraction of all tumor cells that are B cells. Clinical covariates (age at diagnosis, sex, AJCC stage) and overall-survival time/event were both retrieved from TCGAbiolinks. Patients with missing covariates were excluded. Four cohorts (KIRC (ccRCC), LC (LUAD and LUSC), COAD, BRCA) were analyzed separately and OS was right-censored at 3,652 days. Multivariable Cox models, adjusting for B cell abundance, age, sex, and stage, were fit per cohort and hazard ratios (HRs) with 95% CIs were derived.

To assess the prognostic significance of Bregs, transcript-level TPM (RSEM) for TCGA and GTEx samples was downloaded from UCSC Xena (“TcgaTargetGtex_rsem_gene_tpm”). A gene signature of *CD19*, *TGFB1* and *IL10* was scored as the mean TPM across primary-tumor aliquots. Patients were divided into top (Q4) versus bottom (Q1) quartiles. A multivariable Cox proportional-hazards model, adjusting for signature group, age, sex, and AJCC stage yielded hazard ratios (95% confidence intervals).

### Statistical analysis

The distribution of the data was assessed using the Shapiro-Wilk and D’Agostino-Pearson normality tests. For normally distributed and homoscedastic data, statistical significance was determined using a parametric two-tailed *t*-test and one- or two-way analysis of variance (ANOVA). A non-parametric two-tailed Wilcoxon test, Mann-Whitney U test, Kruskal-Wallis test or Brown-Forsythe and Welch ANOVA were applied for non-normally distributed data. For multiple comparisons, the recommended Dunn’s, Tukey’s, Dunnett’s T3, or Šidák’s test was used. The Cox proportional hazard model was used to assess the significance of survival data. No statistical methods were used to determine sample sizes, but our sample sizes are similar to those reported in previous publications^44,57^. All graphs are represented as mean±standard error of the mean (SEM). *p* values are represented as \**p*<0.05, \*\**p*<0.01, \*\*\**p*<0.001, and \*\*\*\**p*<0.0001. Graphing of data and statistical analysis were performed in Prism 10 (GraphPad).

## Notes

https://github.com/IsabellaWithnell/Tissue_specific_B_cell_analysis_pipeline

