## Supplementary Figures for "Tumor-infiltrating CD27^-^IgD^-^ regulatory B cells suppress cytotoxic CD8^+^T cell responses in renal cell carcinoma"

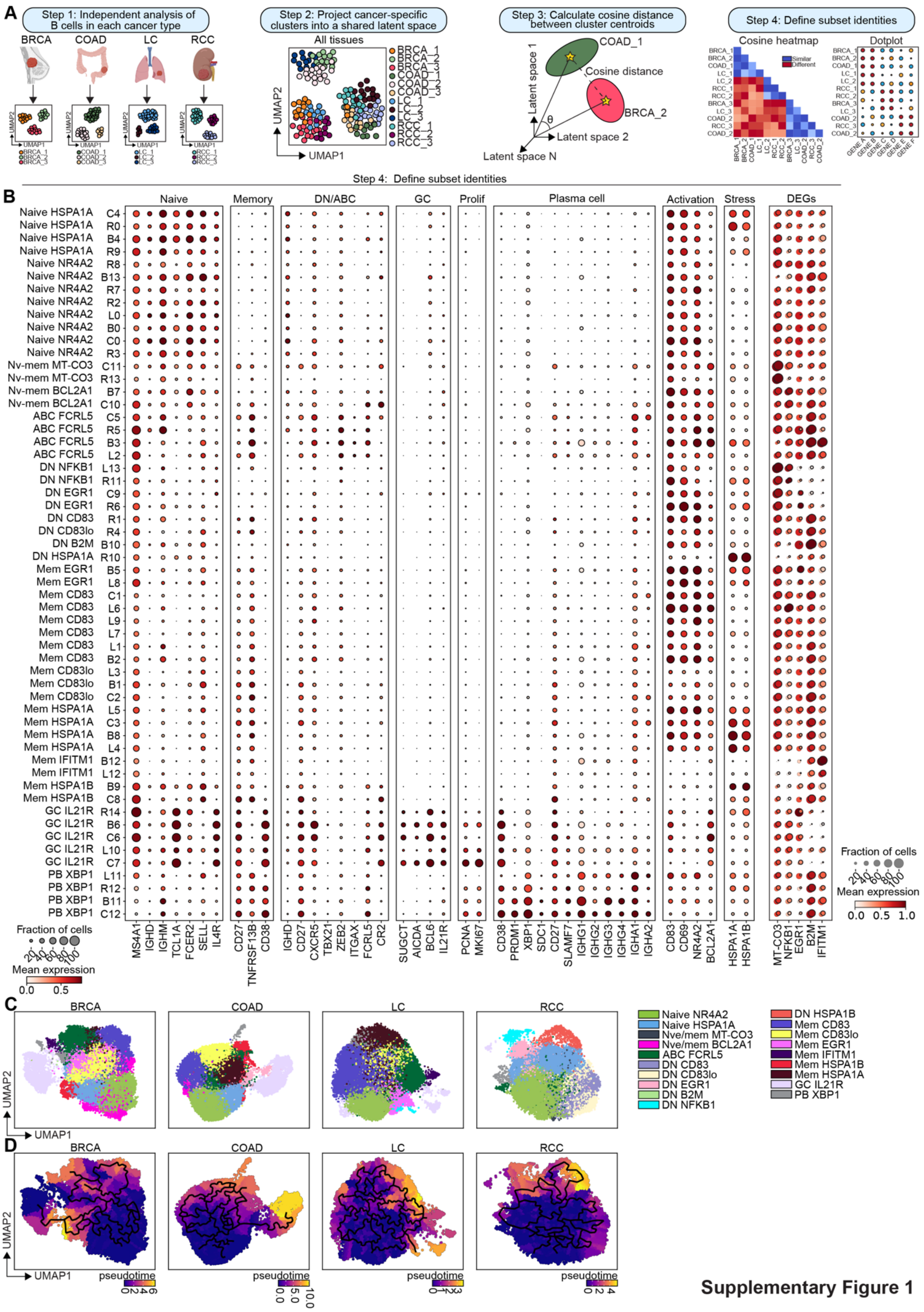


Supplementary Figure 1

1

**Supplementary Figure 1. Cosine distance as a robust approach for grouping similar pan-cancer clusters and assess tissue specificity**

(A) Schematic overview of the cancer-specific bioinformatic analysis pipeline.

(B) Dotplot showing for each cancer-specific B cell cluster the proportion of cells expressing selected key B cell markers (point size) and their mean expression levels (color intensity), min-max scaled across annotated cancer-specific B cell clusters. DN = double negative, mem = memory, ABC = atypical B cell, GC = germinal center, PB = plasmablast, Nve = naive, DEG = differential gene expression, prolif = proliferation.

(C) UMAP visualization of B cell clusters within each cancer-specific latent space for BRCA, COAD, LC, and RCC, annotated according to Figure 1C.

(D) UMAPs showing the inferred pseudotime trajectories within each cancer-specific latent space for BRCA, COAD, LC, and RCC.

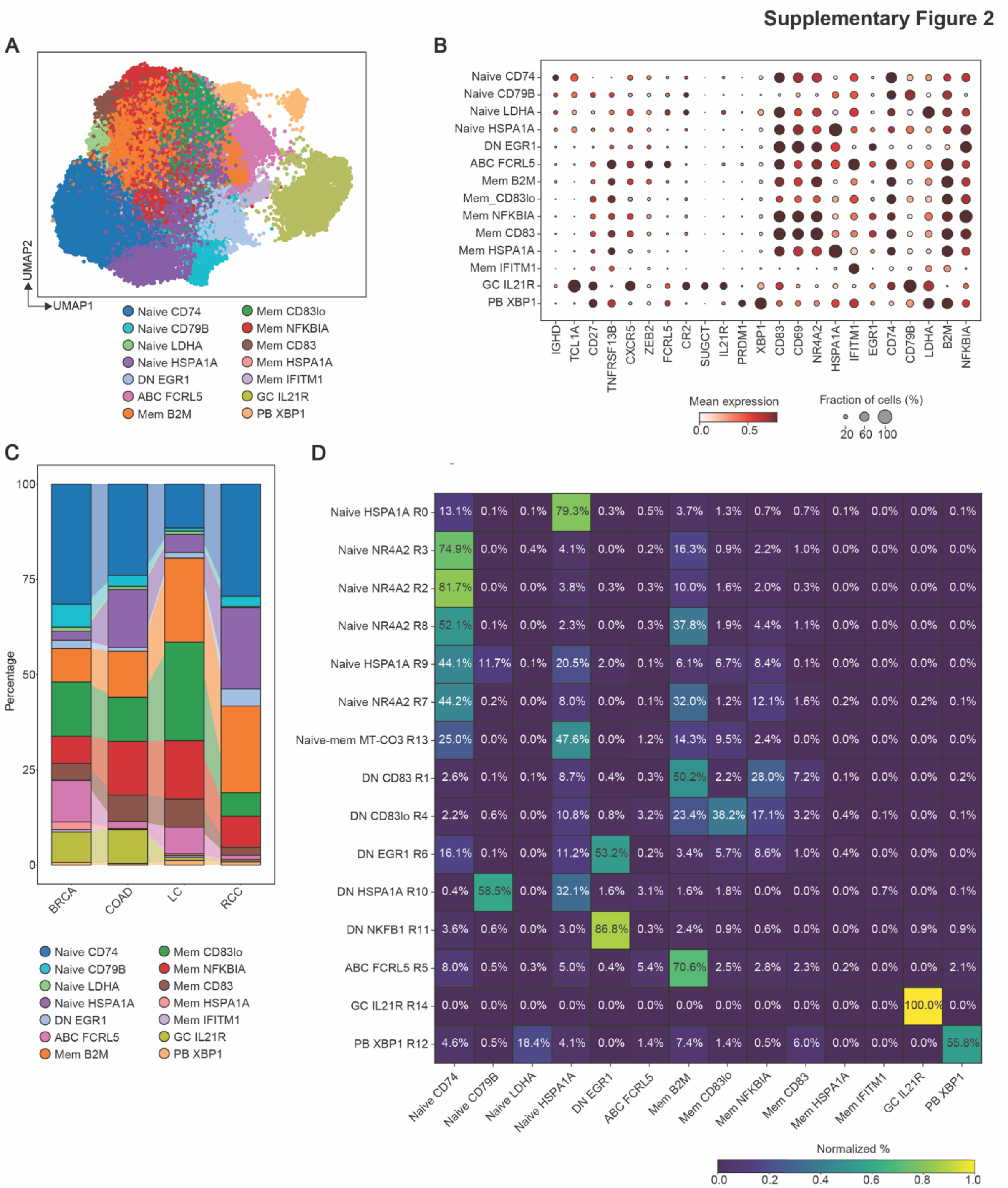


**Supplementary Figure 2. Pan cancer integration and clustering of B cells results in a loss of granularity.**

(A) UMAP showing B cells from BRCA, COAD, LC, and RCC following joint integration and clustering. DN = double negative, mem = memory, ABC = atypical B cell, GC = germinal center, PB = plasmablast.

(B) Dotplot showing the proportion of cells expressing selected markers (point size) and their mean expression levels (color intensity) min-max scaled across B cell clusters from Figure S2A.

(C) Stacked bar chart showing the proportion of annotated B cell clusters in BRCA, COAD, LC, and RCC from the clustering analysis in Figure S2A.

(D) Contingency table showing the overlap between RCC-specific B cell clusters (rows), identified from the cancer-specific analysis, and the clusters defined in the pan-cancer integration (columns, as in Fig. S2A). Percentage indicates the similarity between the RCC-specific clusters in Figure 1 and clusters identified in the pan-cancer analysis in Figure S2A.

**Supplementary Figure 3**

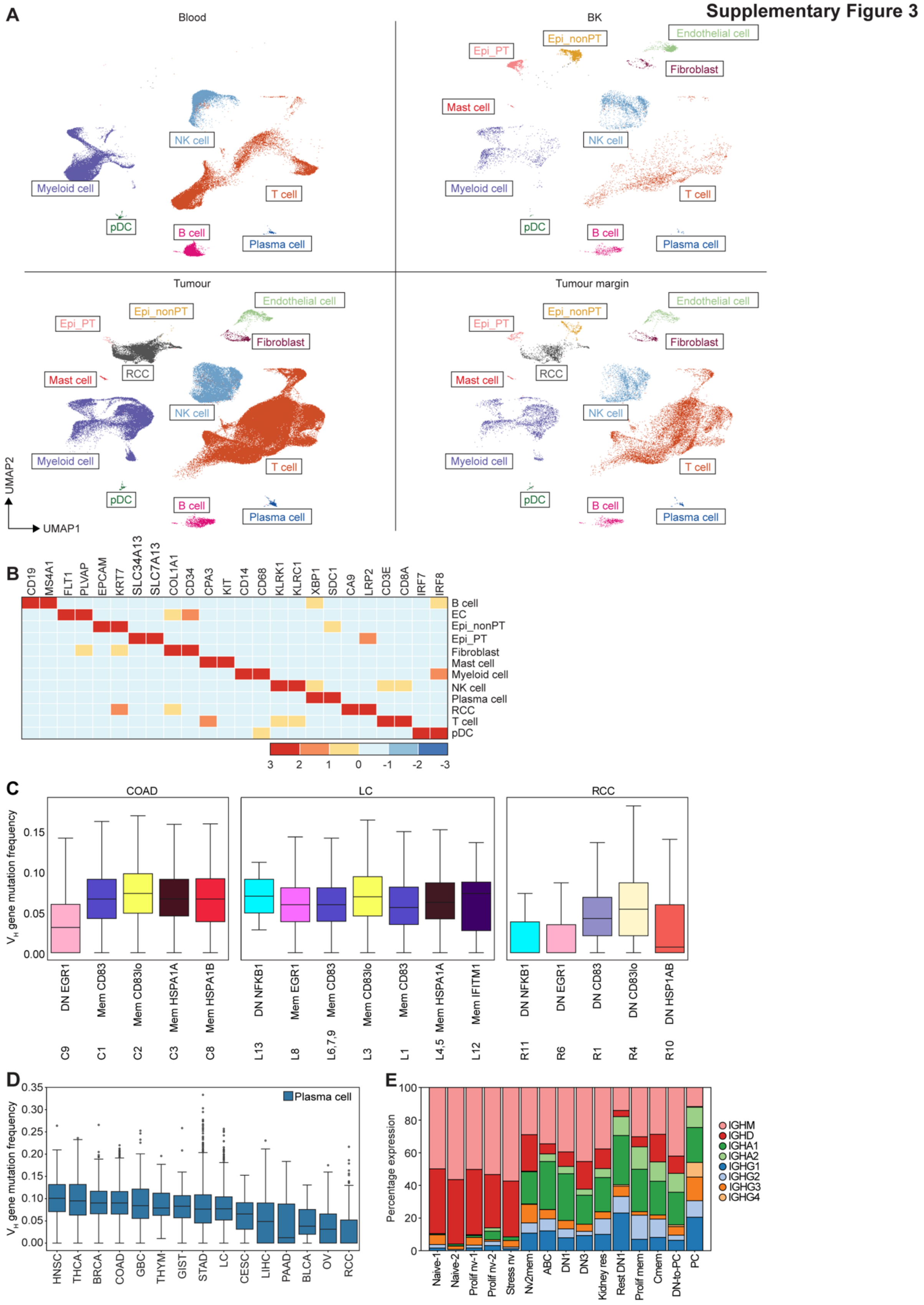


38

**Supplementary Figure 3. Identification of B cell and plasma cell populations in RCC tumor, tumor margin, BK, and peripheral blood.**

(A) UMAPs showing all cell types present in matching tumor, tumor margin, BK, and blood from 10 diagnostically confirmed RCC treatment-naive patients. EC = endothelial cell; Epi_PT = proximal tubule epithelial cell; Epi_nonPT = non-proximal tubule epithelial cell.

(B) Heatmap showing the expression of key marker genes used to define each cell type across paired tissues. EC = endothelial cell; Epi_PT = proximal tubule epithelial cell; Epi_nonPT = non-proximal tubule epithelial cell.

(C) Box plots showing the $V_H$ gene hypermutation frequency in DN B cell and memory B cell clusters across COAD (*n*=7 patients), LC (*n*=8 patients), and RCC (*n*=3 patients) from the cross-cancer analysis in Figure 1 and Figure S1. BRCA excluded due to limited sample size (*n*=1). Data indicate median with interquartile range (IQR), and whiskers indicate minimum and maximum measurement.

(D) Box plots showing the $V_H$ gene hypermutation frequency in previously annotated plasma cells[21] across head and neck squamous cell carcinoma (HNSC), thyroid carcinoma (THCA), BRCA, COAD, gallbladder cancer (GBC), thymoma (THYM), gastrointestinal stromal tumor (GIST), stomach adenocarcinoma (STAD), LC, cervical squamous cell carcinoma and endocervical adenocarcinoma (CESC), liver hepatocellular carcinoma (LIHC), pancreatic adenocarcinoma (PAAD), bladder urothelial carcinoma (BLCA), ovarian serous cystadenocarcinoma (OV), and RCC. Data indicate median with interquartile range(IQR), and whiskers indicate minimum and maximum measurement.

(E) Stacked bar chart showing the expression of immunoglobulin genes across B cell clusters in the tumor, tumor margin, BK and blood from the RCC dataset in Figure 2.

Supplementary Figure 4

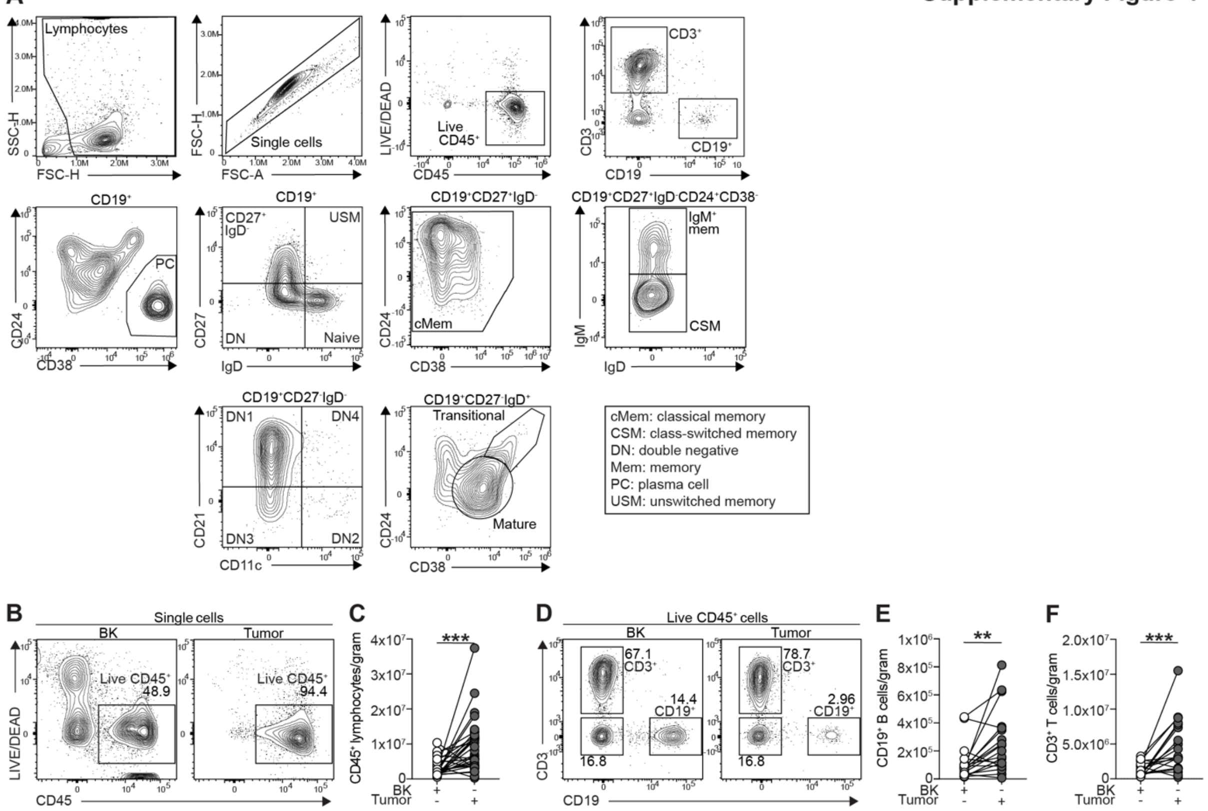


**Supplementary Figure 4. Gating strategy for RCC- and kidney-resident B cell subset identification by flow cytometry and number of B cells per gram of tissue.**

(A) Gating strategy for intratumoral T and B cell identification and for B cell subsets: CD19$^{+}$CD27$^{-}$IgD$^{+}$CD24$^{hi}$CD38$^{hi}$ (transitional), CD19$^{+}$CD27$^{-}$IgD$^{+}$CD24$^{int}$CD38$^{int}$ (mature), CD19$^{+}$CD27$^{-}$IgD$^{+}$ (naive), CD19$^{+}$CD27$^{+}$IgD$^{-}$, CD19$^{+}$CD27$^{+}$IgD$^{-}$CD24$^{+}$CD38$^{-}$ (classical memory (cMem)), CD19$^{+}$CD27$^{+}$IgD$^{-}$CD24$^{+}$CD38$^{-}$IgM$^{-}$ (class-switched memory (CSM)), CD19$^{+}$CD27$^{+}$IgD$^{-}$CD24$^{+}$CD38$^{-}$IgM$^{+}$ (IgM$^{+}$ memory), CD19$^{+}$CD27$^{+}$IgD$^{+}$ (unswitched memory (USM)), CD19$^{+}$CD27$^{-}$IgD$^{-}$ (double negative (DN)), CD19$^{+}$CD27$^{-}$IgD$^{-}$CD21$^{+}$CD11c$^{-}$ (DN1), CD19$^{+}$CD27$^{-}$IgD$^{-}$CD21$^{-}$CD11c$^{+}$ (DN2), CD19$^{+}$CD27$^{-}$IgD$^{-}$CD21$^{-}$CD11c$^{-}$ (DN3), CD19$^{+}$CD27$^{-}$IgD$^{-}$CD21$^{+}$CD11c$^{+}$ (DN4), and CD19$^{+}$CD24$^{lo/-}$CD38$^{hi}$ (plasma cells (PC)).

(B) Representative contour plots showing the frequencies of lymphocytes (CD45$^{+}$ cells) *ex vivo* in BK and matching tumor tissue.

(C) Number of lymphocytes (CD45$^{+}$cells) per gram of BK and matching tumor tissue *ex vivo*; ***$p$<0.001. $n$=27 biologically independent samples. Paired two-tailed Wilcoxon test.

(D) Representative contour plots showing the frequencies of CD19$^{+}$ B cells and CD3$^{+}$ T cells *ex vivo* in BK and matching tumor tissue.

(E) Number of CD19$^{+}$B cells per gram of BK and matching tumor tissue *ex vivo*; **$p$<0.01. $n$=17 biologically independent samples. Paired two-tailed Wilcoxon test

(F) Number of $CD3^{+}$T cells per gram of BK and matching tumor tissue *ex vivo*; ***$p<0.001$. $n=18$ biologically independent samples. Paired two-tailed Wilcoxon test.

**Supplementary Figure 5**

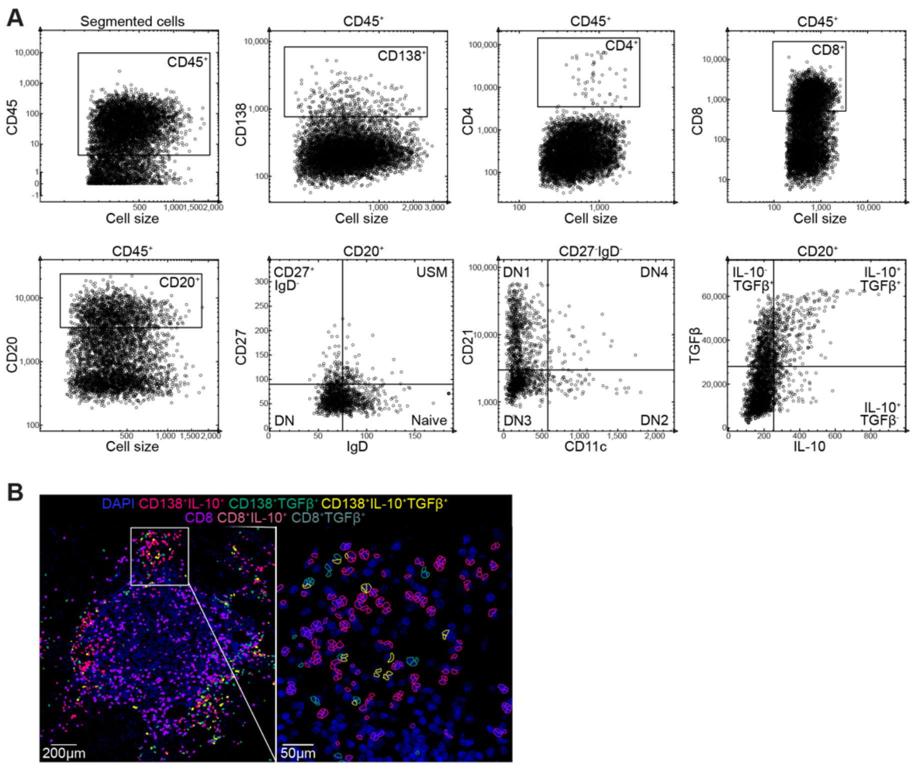


**Supplementary Figure 5. Gating strategy for RCC-resident B cell subsets by MACSima Imaging Cyclic Staining (MICS), and proximity of CD138$^{+}$Bregs to CD8$^{+}$T cells.**

(A) Gating strategy for intratumoral CD4$^{+}$ T cell, CD8$^{+}$ T cell, CD20$^{+}$ B cell, and CD138$^{+}$ plasma cell identification and for B cell subsets: CD20$^{+}$CD27$^{-}$IgD$^{-}$ (double negative (DN)), CD20$^{+}$CD27$^{+}$IgD$^{-}$, CD20$^{+}$CD27$^{+}$IgD$^{+}$ (unswitched memory (USM)), CD20$^{+}$CD27$^{-}$IgD$^{+}$ (naive), CD20$^{+}$CD27$^{+}$IgD$^{-}$, CD20$^{+}$CD27$^{-}$IgD$^{-}$CD21$^{+}$CD11c$^{-}$ (DN1), CD20$^{+}$CD27$^{-}$IgD$^{-}$CD21$^{-}$CD11c$^{+}$ (DN2), CD20$^{+}$CD27$^{-}$IgD$^{-}$CD21$^{-}$CD11c$^{-}$ (DN3), CD20$^{+}$CD27$^{-}$IgD$^{-}$CD21$^{+}$CD11c$^{+}$ (DN4), CD20$^{+}$IL-10$^{+}$TGFβ$^{-}$, CD20$^{+}$IL-10$^{-}$TGFβ$^{+}$, and CD20$^{+}$IL-10$^{+}$TGFβ$^{+}$ B cells.

(B) Spatial analysis of CD138$^{+}$ PC Bregs and CD8$^{+}$ T cells in the TLS. Staining of DAPI (dark blue) overlaid with cell segmentation (gating strategy shown in Figure S5A) of CD138$^{+}$IL-10$^{+}$ Bregs (dark pink), CD138$^{+}$TGFβ$^{+}$Bregs (green), CD138$^{+}$IL-10$^{+}$TGFβ$^{+}$Bregs (yellow), CD8$^{+}$T cells (purple), CD8$^{+}$IL-10$^{+}$T cells (light pink) and CD8$^{+}$TGFβ$^{+}$T cells (grey). Scale bar = 200µm (Panel 1), 50µm for (Panel 2).

**Supplementary Figure 6**

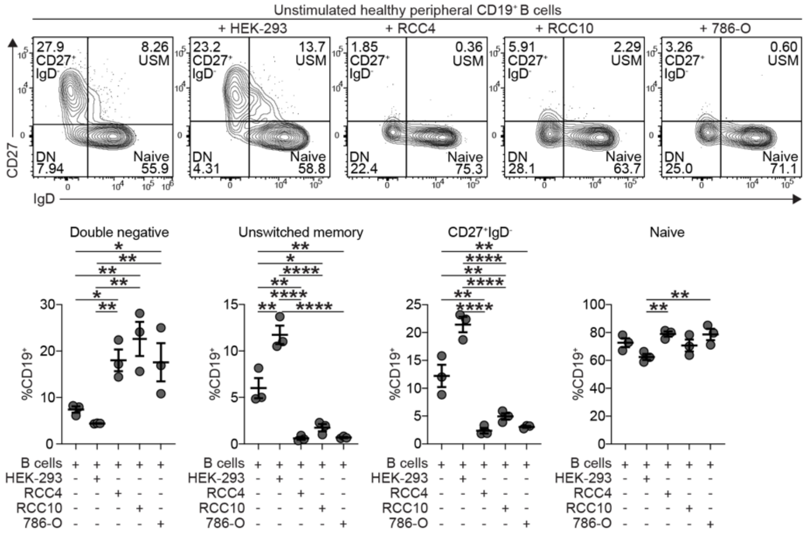


**Supplementary Figure 6. RCC cell lines convert circulating healthy B cells into DN B cells.**

Representative contour plots and cumulative data showing the frequencies of $CD19^+CD27^-IgD^-$ (double negative (DN)), $CD19^+CD27^+IgD^+$ (unswitched memory (USM)), $CD19^+CD27^+IgD^-$, and $CD19^+CD27^-IgD^+$ (naive) B cells after co-culture of healthy peripheral blood B cells alone or with HEK-293, RCC4, RCC10, or 786-O cell lines for 72h; **p*<0.05, ***p*<0.01, ****p*<0.001, *****p*<0.0001. *n*=3 biologically independent samples. One-way ANOVA with Tukey’s test for multiple comparisons. Error bars represented as mean±SEM.

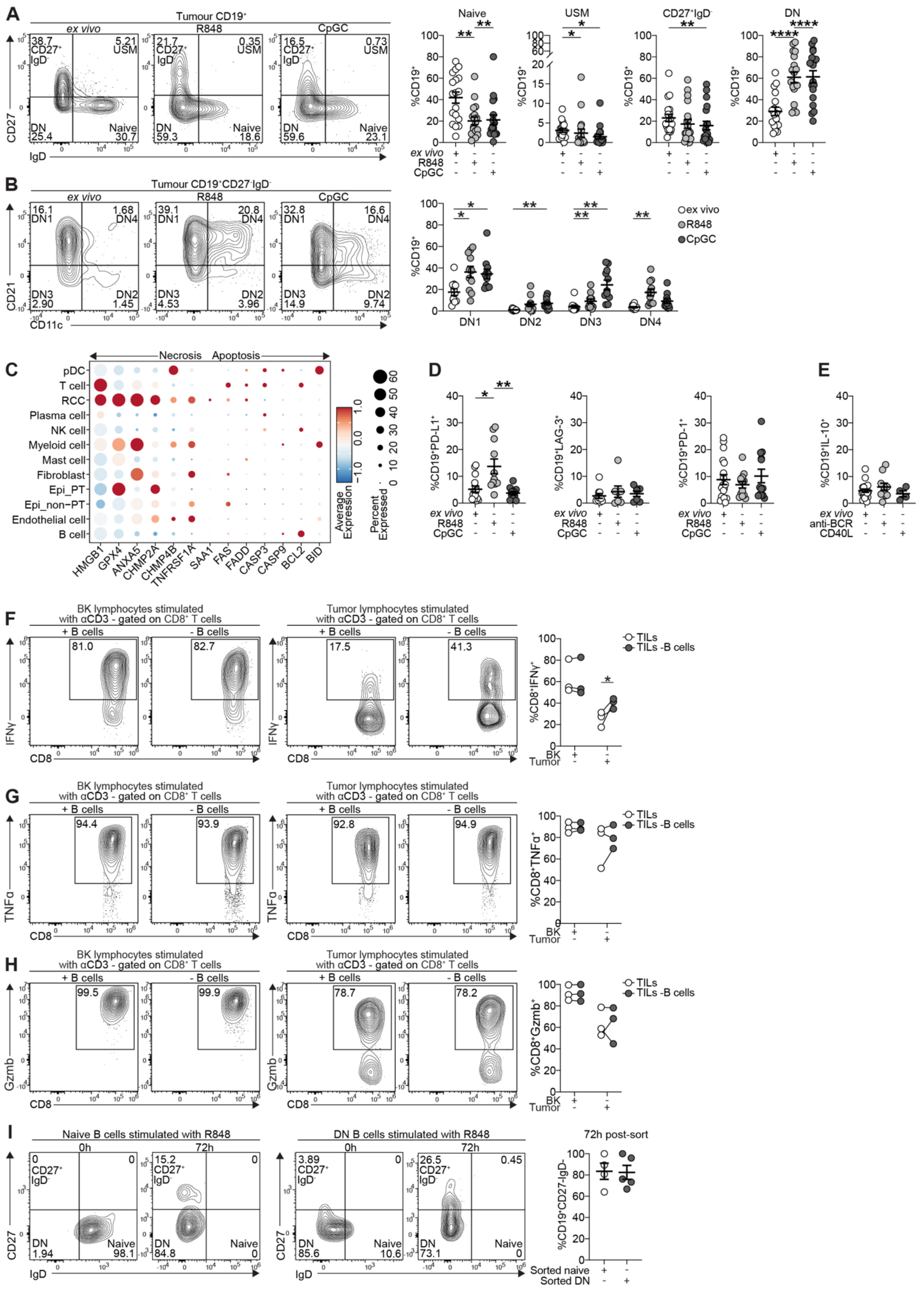


127

**Supplementary Figure 7**

**Supplementary Figure 7. R848 and CpGC induce the differentiation of DN1 B cells, but fail to expand LAG-3$^+$ or PD-1$^+$ intratumoral B cells; B cell depletion from tumor, but not from BK, partially rescues intratumoural IFNγ$^+$CD8$^+$ T cells frequency.**

(A) Representative contour plots and cumulative data showing the frequencies of CD19$^+$CD27$^-$IgD$^+$ (naive), CD19$^+$CD27$^+$IgD$^+$ (unswitched memory (USM)), CD19$^+$CD27$^+$IgD$^-$, and CD19$^+$CD27$^-$IgD$^-$ (double negative (DN)) B cells, *ex vivo* and following 72h stimulation with R848 or CpGC; *$p<0.05$, **$p<0.01$, ****$p<0.0001$. $n$=18 biologically independent samples. Naive, USM, and CD27$^+$IgD$^-$ analyzed by Fridman test with Dunn's test for multiple comparisons, DN analyzed by one-way ANOVA with Tukey's test for multiple comparisons. Error bars represented as mean±SEM.

(B) Representative contour plots and cumulative data showing the frequencies of CD19$^+$CD27$^-$IgD$^-$CD21$^+$CD11c$^-$ (DN1), CD19$^+$CD27$^-$IgD$^-$CD21$^-$CD11c$^+$ (DN2), CD19$^+$CD27$^-$IgD$^-$CD21$^-$CD11c$^-$ (DN3) and CD19$^+$ CD27$^-$IgD$^-$CD21$^+$CD11c$^+$ (DN4) B cells, *ex vivo* and following 72h stimulation with R848 or CpGC; *$p<0.05$, **$p<0.01$. *Ex vivo* $n$=12, R848 and CpGC $n$=11 biologically independent samples. Two-way ANOVA with Tukey's test for multiple comparisons. Error bars represented as mean±SEM.

(C) Dotplot showing the proportion of cells expressing genes associated with necrosis and apoptosis cell death pathways (dot size), and their average expression levels (color intensity) within RCC-resident cells.

(D) Cumulative data showing PD-L1, LAG-3, and PD-1 expression by tumor-infiltrating CD19$^+$ B cells *ex vivo* (white) and following 72h stimulation with R848 (light grey) or CpGC (dark grey). PD-L1 *ex vivo* $n$=15, R848 and CpGC $n$=22; for LAG-3 *ex vivo* $n$=10, R848 and CpGC $n$=8; for PD-1 *ex vivo* $n$=18, R848 and CpGC $n$=13 biologically independent samples. Kruskal-Wallis test with Dunn's test for multiple comparisons. Error bars represented as mean±SEM.

(E) Cumulative data showing IL-10 expression by tumor-infiltrating CD19$^+$ B cells *ex vivo* (white) and following 72h stimulation with anti-IgM/IgG/IgA (anti-BCR) (light grey) or CD40 ligand (CD40L) (dark grey). For *ex vivo* $n$=26, anti-BCR $n$=9, and CD40L $n$=4 biologically independent samples. Mixed-effects one-way ANOVA with Dunnett's test for multiple comparisons. Error bars represented as mean±SEM.

(F) Representative contour plots and cumulative data showing the frequencies of anti-CD3-stimulated IFNγ$^+$CD8$^+$ T cells from the BK and tumor, in the presence or absence of B cells; *$p<0.05$. $n$=3 biologically independent samples. Two-way ANOVA with Šidák's test for multiple comparisons.

(G) Representative contour plots and cumulative data showing the frequencies of anti-CD3-stimulated TNFα$^+$CD8$^+$ T cells from the BK and tumor, in the presence or absence of B cells. $n$=3 biologically independent samples. Two-way ANOVA with Šidák's test for multiple comparisons.

(H) Representative contour plots and cumulative data showing the frequencies of anti-CD3-stimulated Gzmb$^+$CD8$^+$ T cells from the BK and tumor, in the presence or absence of B cells. *n*=3 biologically independent samples. Two-way ANOVA with Šidák's test for multiple comparisons.

(I) Representative contour plots and cumulative data showing the frequencies of naive and DN B cells at the time of FACS-sorting (0h) and following a 72h *in vitro* culture with R848. *n*=5 independent patient samples. Error bars represented as mean±SEM.
